# Discovery of an APP-Selective BACE1 Inhibitor for Alzheimer’s Disease

**DOI:** 10.1101/2023.08.04.552030

**Authors:** Jesus Campagna, Barbara Jagodzinska, Dongwook Wi, Chunni Zhu, Jessica Lee, Whitaker Cohn, Michael Jun, Chris Elias, Olivier Descamps, Qiang Zhang, Olivia Gorostiza, Karen Poksay, Patricia Spilman, Dale Bredesen, Varghese John

## Abstract

Inhibition of amyloid precursor protein (APP) beta-site cleaving enzyme 1 (BACE1; BACE) has been a target for Alzheimer’s disease (AD) therapeutic development, but has been impaired by off-target effects of clinically evaluated inhibitors, including inhibition of cleavage of non-APP substrates. Here, we report our identification of a BACE inhibitors series that are not only selective for the APP substrate, but also for BACE1 as the targeted enzyme. These APP-selective fluoro aminohydantoin (FAH) inhibitor compounds were identified by screening a compound library for inhibition of BACE cleavage of a maltose binding protein (MBP)-conjugated-APPC125 substrate followed by IC50 determination using the P5-P5’ substrate assay. In multiple substrate and enzyme cell-free assays, the lead compound FAH65 displayed substrate selectivity for inhibition of APP cleavage, with little activity against BACE substrates neuregulin 1 (NRG1) or p-selectin glycoprotein ligand -1 (PSGL1). We also demonstrate FAH65 shows little inhibitory activity against the enzyme cathepsin D (Cat D) or BACE2. FAH65 inhibits production of BACE cleavage products soluble APPβ (sAPPβ) and the β C-terminal fragment (βCTF), as well as amyloid-β (Aβ)1-40 and 1-42, *in vitro* in cells and *in vivo* in an animal model of AD. In a murine model of AD, FAH65 improved the discrimination score in the Novel Object Recognition (NOR) memory testing paradigm. The active enantiomer of FAH65, FAH65E(-), was obtained and tested in in vivo pharmacokinetic and pharmacodynamic (PK/PD) analysis, wherein it displayed good brain-penetrance and target engagement. Given its demonstrated selectivity for both enzyme and substrate, along with evidence it can improve cognitive performance in an animal model, FAH65 and its E(-) enantiomer merit continued pre-clinical development towards clinical testing as an APP-selective BACE1 inhibitor. Such a candidate would reduce Aβ levels and overcome the deleterious effects of the non-selective BACE1 inhibitors that have failed in the clinic and potentially could be used as a maintenance therapy along with or following clearance of Aβ from the brain with the approved antibody therapy for AD.

## Introduction

Brain tissue of Alzheimer’s disease (AD) patients is characterized by the presence of neuritic plaques largely composed of amyloid-β (A*β*) (1, 2) and neurofibrillary tangles resulting from the hyperphosphorylation of the protein tau (3-5). A*β* is the product of step-wise cleavage of amyloid precursor protein (APP) by the *β*-site APP–cleaving enzyme 1 (BACE-1; BACE) and *γ*-secretase (6, 7). In an alternative cleavage pathway, α-secretase (putatively ADAM10) cleavage of APP results in production of trophic, synapse-supporting peptides soluble APPα (sAPPα) and the α C-terminal fragment (αCTF)(8-10).

While the amyloid cascade hypothesis of AD has come under scrutiny in the last decade due to the failure of Aβ- and BACE-directed potential therapies in the clinic (11), it is supported by the genetics underlying familial forms of the disease that enhance amyloid production (12, 13), by identification of gene variants that confer protection against cognitive decline (14); and by recent clinical findings that the amyloid-directed antibodies (Aduhelm^®^), lecanemab (Lequembi^®^), and donanemab slow cognitive decline in early AD (15-18).

Antibody-based biologic therapies present challenges as feasible therapeutics due to their limited brain-permeability, high cost, and requirement for IV infusion at hospitals or infusion centers (19). An orally-available, brain-permeable small-molecule therapeutic that inhibits BACE cleavage of APP would overcome those limitations and would not only decrease amyloid plaque formation, but also the upstream formation of the Aβ oligomers that are implicated in synaptic loss and cognitive decline (11, 20).

The discovery that an A673T mutation (Icelandic) in APP at the P2’ residue from the BACE cleavage site, reduces BACE cleavage of APP, protects against both AD and other age-related cognitive decline (14) provides further strong support for the hypothesis that Aβ production, and that BACE cleavage specifically, is a critical event in AD pathogenesis. Thus, inhibition of BACE as a strategy for therapeutic development for AD has been a focus of many research groups.

A critical limitation of BACE inhibitory strategies is the potential inhibition of cleavage of non-APP substrates, resulting in side effects. Inhibition of non-APP substrates such as neuregulin 1 (NRG1), p-selectin glycoprotein ligand 1 (PSGL1) and gp-130 raise concern (21-25). Therefore, the optimal BACE inhibitor would be one that would inhibit the cleavage of APP selectively. Such a therapeutic would represent a new class of AD therapeutics: APP-selective BACE inhibitors (ASBIs). ASBIs would potentially selectively inhibit APP binding to the BACE active site compared to other BACE substrates resulting in potent inhibition of APP cleavage by BACE compared to inhibition of enzyme cleavage of other substrates, as previously described (26).

Development of an ASBI may not only provide a potential new treatment for AD and the condition that often precedes it, Mild Cognitive Impairment (MCI) due to AD, but may do so with reduced off-target effects as compared to BACE inhibitors that have been clinically tested and are not selective for APP.

Our strategy to discover ASBIs is to use a larger APP fragment using a maltose binding protein (MBP)-conjugated-APPC125 substrate to screen for BACE1 inhibitors. As reported in Descamps et al. (26), by screening a clinical compound library, we previously identified flavonoids that acted as ASBIs in cell models and one (galangin) that showed increased brain levels when delivered to mice as a pro-drug.

Here we describe our identification of the screening hit phenytoin as a weak inhibitor of BACE cleavage of the MBP-APPC125 substrate and, through medicinal chemistry optimization efforts that led to an APP selective fluoro aminohydantoin (FAH) series, and the discovery of a highly potent and selective lead compound FAH65 that shows potent inhibition of BACE in vitro and in vivo, along with improvement in memory in an AD mouse model after oral treatment.

## RESULTS

### Screening and analog generation by medicinal chemistry identifies lead FAH65

We identified a known anti-epilepsy hydantoin drug, phenytoin, as a screening hit in the MBP-APPC125 BACE cleavage screening assay [26]. Initial exploratory medicinal chemistry on the phenyl rings (A and B-rings) of the hydantoin hit revealed no significant improvement in BACE inhibition. Further modification of the hydantoin scaffold to an aminohydantoin led to a micromolar BACE inhibitor fluoroaminohydantoin, FAH1, with A-ring fluorine substitutions that inhibited Aβ42 in cells and was brain permeable. Further modification of the B-ring led to a more potent low micromolar inhibitor, FAH3, that also inhibited Aβ in cell models and had brain permeability, but also showed selectivity for APP cleavage by BACE versus other substrates such as NRG1. The BACE IC50s in the cell-free P5-P5’ activity assay for FAH1 and FAH3 were ∼10 μM and ∼1-3 μM, respectively (See **Supplementary Figure S1**).

Interestingly, removal of a fluorine from the A-ring led to a more potent analog, FAH17 (IC50 ∼ 0.3-1μM) and further replacement of the fluorine with a pyrimidine ring gave lead compound FAH65 (IC50 ∼ 0.01-0.02 μM) (**Figure 1**). Details of the structure-activity strategy will be described in a future manuscript.

**Figure 1.**
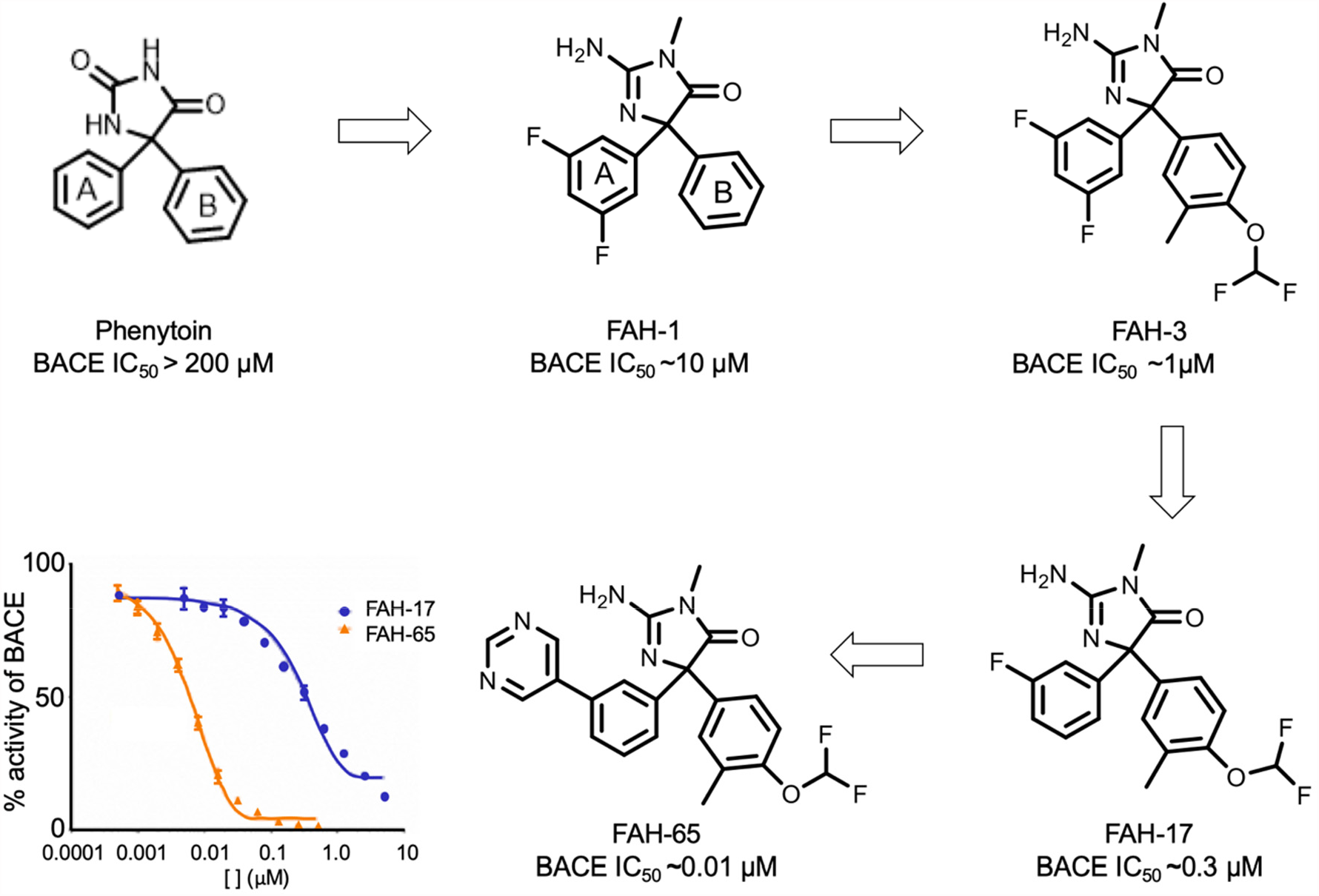
Analog structures and inhibition of BACE. The structures of phenytoin, FAH1, FAH3, FAH17 and FAH65 are shown, as well as the dose-response curve for FAH17 and FAH65 in the P5-P5’ BACE inhibition assay.

### FAH65 selectively targets BACE and inhibition of APP cleavage

Ongoing selectivity monitoring using a SEAP-neuregulin 1 (NRG1) assay demonstrated the analogs were selective and did not inhibit BACE cleavage of NRG1. Specifically, neither FAH17 nor FAH65 show potency in cleavage of either NRG1 or PSGL1 (both EC_50_s > 10 µM) vs. BACE-APP (IC_50_s of 0.3 µM and 0.01 µM, respectively) in cell-free assays as compared to the inhibitor BACEIV that has little selectivity for APP (BACE-APP IC_50_ ∼0.04 µM) vs NRG1 or PSGL1 (both EC_50_ ∼0.05 µM) (**Fig. 2A and B, respectively**). In comparison, clinically-studied BACE inhibitors verubecestat (veru; MK-8931) and lanabecestat (lana; AZD3293) show no selectivity for APP (IC_50_s of 0.002 and 0.004 µM, respectively) vs. NRG1 (IC_50_s of < 0.01 µM for both); with substrate selectivity: NRG1/APP IC_50_ of >200 for FAH65 and <2.5 for both veru and lana (**Supplementary Table S1**).

**Figure 2.**
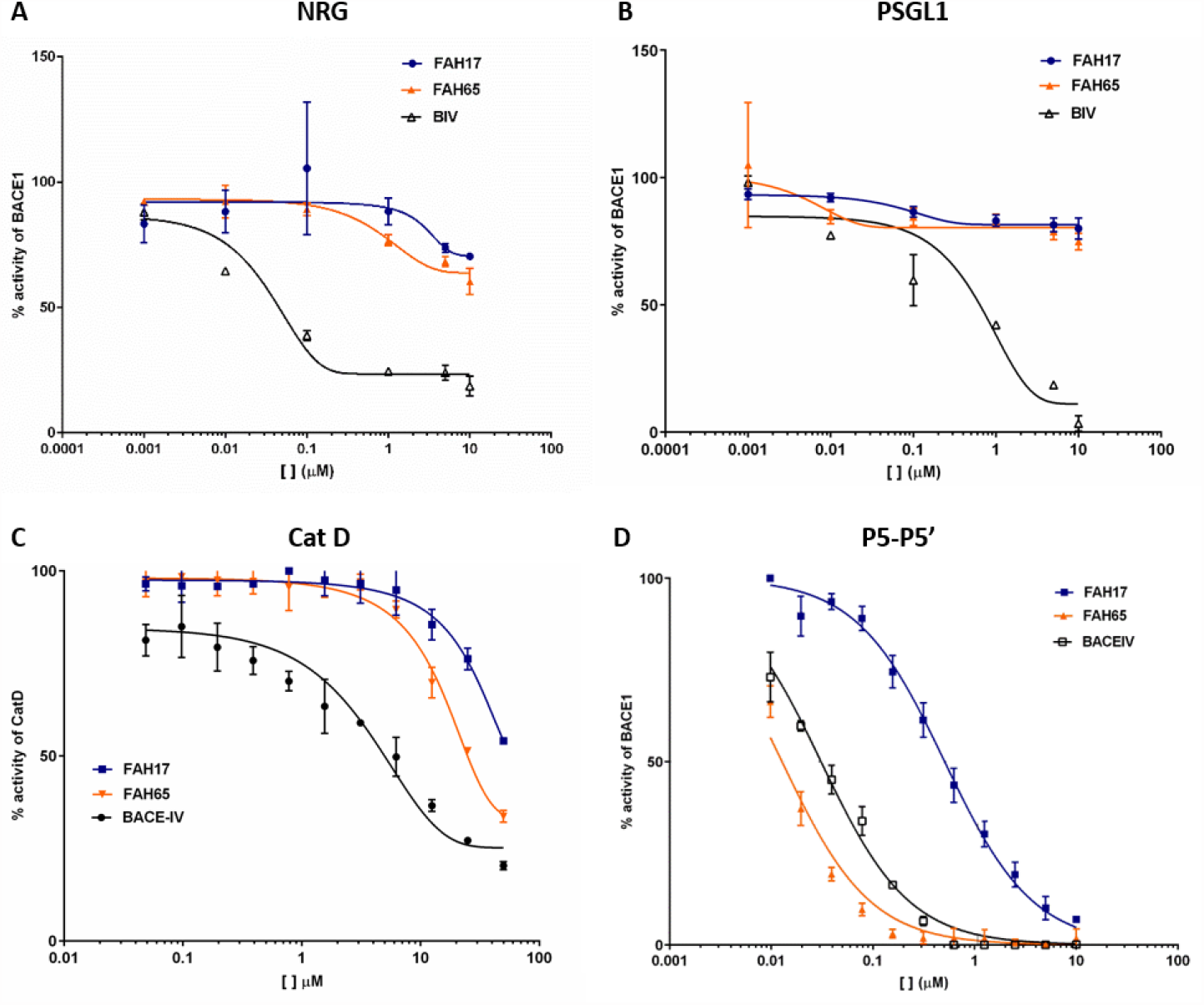
FAH65 selectivity. Dose-response curves for FAH17 and FAH65 as compared to BACE1 inhibitor BACEIV are shown for the substrates (A) NRG1 and (B) PSGL1 in cell-free assays. Dose-response curves for inhibit of (C) cathepsin D (cat D) and (D) BACE in the P5-P5’ assay are shown.

FAH17 and FAH65 were also found to be selective for BACE as the target enzyme. Neither analog inhibited cathepsin D (CAT D) activity (**Fig. 2C**), and lead FAH65 did not inhibit BACE2 activity (**Supplementary Fig S2**); whereas both inhibit P5-P5’ cleavage by BACE 1 (**Fig. 2D**). The FAH65 IC_50_s for Cat D, BACE2, and BACE are ∼47 µM, >1 μM and ∼0.007 µM, respectively.

### FAH65 inhibits sAPPβ and Aβ1-42 production in APP-expressing cells in vitro

Both FAH17 and FAH65 display the ability to decrease production of BACE cleavage product soluble APPβ (sAPPβ) and amyloid-β 1-42 (Aβ1-42) in Chinese hamster ovary cells expressing wildtype human APP (CHO-7W) (**Fig. 3A and B**, respectively), but only FAH65 significantly decreases Aβ1-42 in CHO cells that express APP with the Swedish (Swe) K595N/M596L mutation (**Fig. 3C**). Cells expressing APP with the Swedish mutation were tested in anticipation of in vivo studies in AD model mice that express human APP with the Swe mutation.

**Figure 3.**
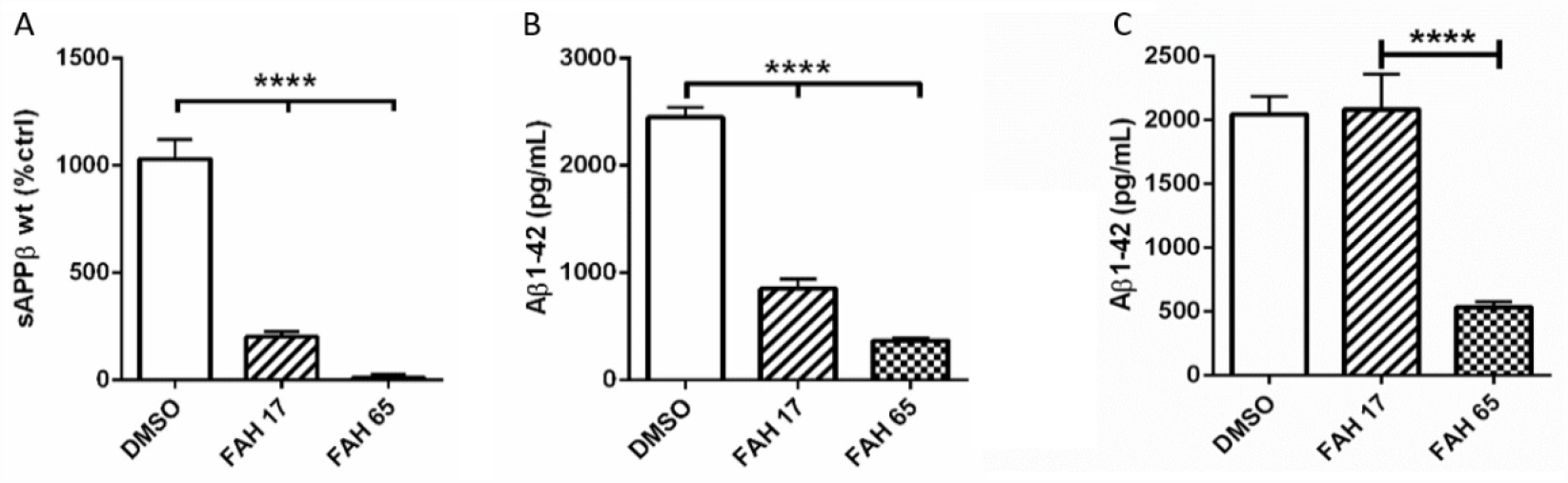
sAPPβ and Aβ1-42 production in CHO-7W and CHO-Swe cells. Levels of sAPPβ (A) and (B) Aβ1-42 in CHO-7W wildtype APP, and (C) Aβ1-42 in APP Swe cell lysates after 24-hour treatment at 1 μM are shown. Data graphed as the mean and SEM. Statistical analysis performed using one-way ANOVA and Dunnett’s post-hoc analysis.

### Pilot APP binding experiments of the FAH analogs by surface plasmon resonance (SPR)

The interaction of FAH3,17 and 65 were evaluated using the SPR protocol described previously in Descamps et al., (26). The FAHs showed varying interaction with APP, when compared to the BACE-IV inhibitor (**Supplementary Table S3)**. The binding to APP may contribute to the selectivity seen with the FAHs for BACE cleavage of APP substrate relative to NRG1 and PSGL1.

### FAH65 displays favorable physiochemical properties and brain permeability in mice

FAH65 was determined to have kinetic solubility at 88 μM in water, plasma stability at t_1/2_ > 180 min. and, in the Parallel Artificial Membrane Permeability (PAMPA) assay, a Pm of 1.09, predicting it would be brain-permeable. Its half-life in liver microsomes is ∼38 min and it binds human serum albumin (HSA) with an free unbound (F_u_) of 0.44%. In brain tissue the F_u_ was 3.1% for (see **Supplementary Methods** and **Supplementary Table S1)**. In hERG testing, FAH65 showed potassium channel binding below 50%, ranging from 38.7-41% at 10uM (**see Supplementary Methods**).

After SQ delivery of FAH65 to mice at 10 or 30 mg/kg, compound levels over 3000 ng/mL were observed in plasma (**Fig. 4A**). After oral delivery of FAH65 at 30 mg/kg, the plasma C_max_ of ∼50 ng/mL was seen at 2 hours and the brain C_max_ of 74 ng/g at 1 hour (**Fig. 4B**).

**Figure 4.**
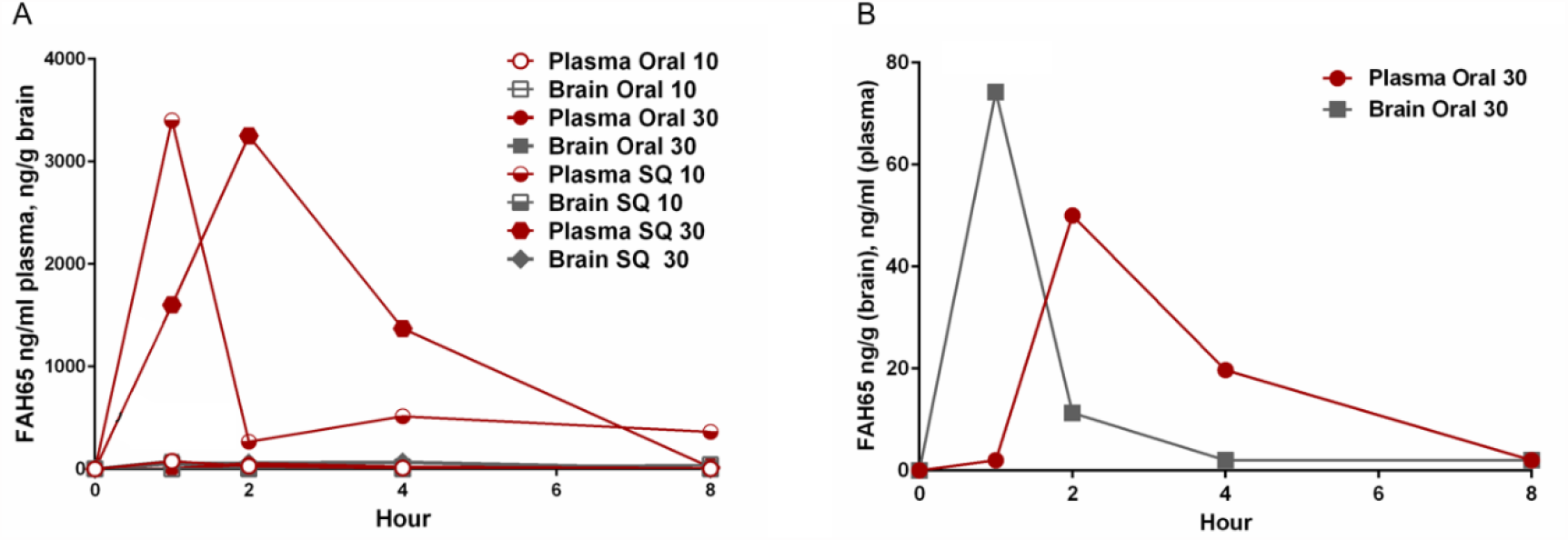
Pharmacokinetics of FAH65 in mice. (A) Plasma and brain levels of FAH65 at 10 and 30 mg/kg 1, 2, 4 and 8 hours after delivery by either the oral or subcutaneous (SQ) routes are shown. (B) Plasma and brain levels of FAH65 at 30 mg/kg after oral delivery are shown.

### Pilot PK/PD comparison of FAH65 and verubecestat (veru) in rats

A pilot PK/pharmacodynamic (PD) study was performed in rats to compare FAH65 with known BACE inhibitor veru [27]. After oral gavage delivery at 30 mg/Kg, the brain FAH65 C_max_ of 182 ng/g (**Supplementary Fig. S3A**) and veru C_max_ of 157 ng/ng (**Supplementary Fig. S3B**) were observed at 2 hours post-delivery, showing similar brain permeability. In cerebral spinal fluid (CSF), APP BACE cleavage product sAPPβ was the lowest at 2 hours post-delivery of veru and 3 hours post-delivery for FAH65 (1 and 2 hours samples were not available for assay), and were lower with FAH65 (**Supplementary Fig. S3C**). In brain (medial cortex), sAPPβ was lowest 6 hours post-delivery of FAH65 (**Supplementary Fig. S3D**). In addition to sAPPβ, BACE cleavage of APP produces the β C-terminal fragment (βCTF), which can then be cleaved by γ-secretase to produce Aβ1-40 and/or Aβ1-42. Aβ1-40 was also lower in CSF from FAH65-treated as compared with veru-treated rats (**Supplementary Fig. S3E**). Aβ1-42 in brain varied with FAH65 and veru treatment (**Supplementary Fig. S3F**). Inhibition of APP cleavage by BACE likely provides more available substrate for α-secretase cleavage and sAPPα generation; sAPPα levels in brain medial cortex were higher with FAH65 (**Supplementary Fig. S3G**).

### FAH65 improved cognitive performance in vivo in a murine model of AD

In the pilot study (n = 8) of oral 30 mkd FAH65 for 10 days in B254 mice, target engagement in the brain was evidenced by the significant decrease in APP β-CTF in brain tissue of FAH65-, as compared to vehicle-, treated mice (**Fig. 5A**). While the means for Aβ 1-42 and the p-tau/t-tau ratio were lower in FAH65 vs. vehicle-treated mice, the differences were not significant (**Fig. 5B and C)**. The performance of FAH65-treated mice in the Novel Object Recognition (NOR) testing paradigm (28) was not significantly increased as compared to vehicle-treated mice (**Fig. 5D**). Two hours after the last dose of FAH65 on the last day of treatment (time of euthanasia), FAH65 showed modest brain penetrance, with a mean of ∼ 20 ng/g in brain (**Fig. 5F**).

**Figure 5.**
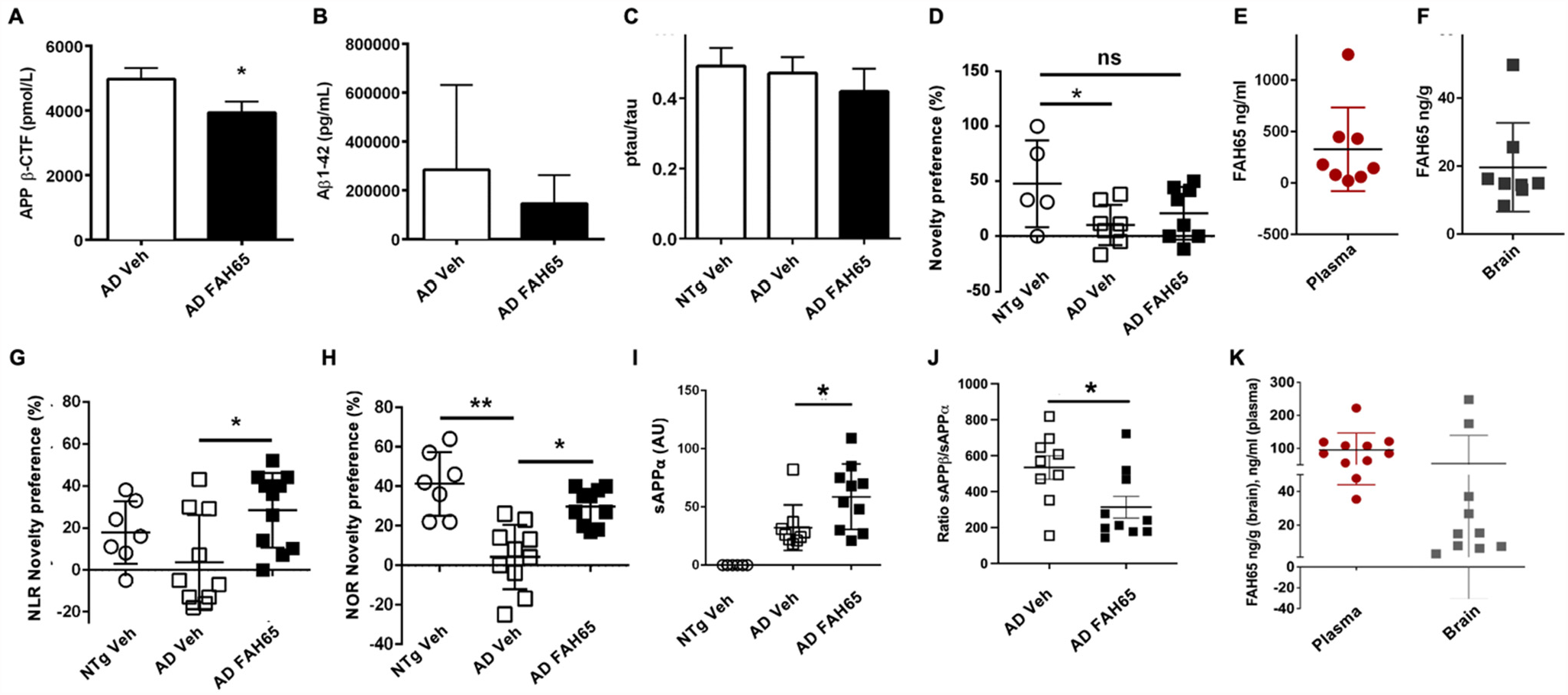
FAH65 efficacy in AD model mice. Pilot study brain levels of (A) APP β-CTF, (B) Aβ 1-42, and (C) the p-tau/total tau ratio; as well as (D) novelty preference in the NOR testing paradigm, (E) plasma and (F) brain levels of FAH65 in non-transgenic (NTg; n = 5), AD model vehicle-treated (AD Veh; n = 8), and FAH65-treated (AD FAH65; n = 8) AD model mice are shown. Follow-up study (G) novelty preference in NLR, and (H) NOR are shown, as well as brain levels of (I) sAPPα, and (J) the sAPPα/sAPPβ ratio. (K) Plasma and brain levels of FAH65 are shown. NTg Veh (n = 7), AD Veh (n = 9), and AD FAH65 (n = 11) mice. Data graphed as the mean and SEM. Statistical analysis performed using one-way ANOVA with Dunnett’s group comparisons; *p < .05 and **p ≤ .01.

In the follow-up study (n = 11) of oral 30 mkd FAH65 for 26 days, a significant increase in novelty preference was observed in both NLR and NOR for FAH65-vs. vehicle-treated mice (**Fig. 5G and H**). Both sAPPα (Fig. 5I), which may be increased by BACE inhibition, and the sAPPβ/sAPPα ratio (**Fig. 5J**) were significantly lower in brain tissue of FAH65-as compared to vehicle-treated mice supporting target engagement. The mean brain level of FAH65 was ∼50 ng/g 2 hours after dosing (**Fig. 5K**).

### Separation and testing of the active enantiomer of FAH65

The (+) and (-) enantiomers of FAH65 were separated using supercritical fluid chromatography as described in **Supplementary Methods** and underwent assessment for relative BACE inhibition. In the P5-P5’ assay, the (-) enantiomer of FAH65, FAH65E (-), shows high activity (IC_50_ ∼0.005 µM), whereas the FAH65 (+) enantiomer is inactive (**Fig. 6A**). FAH65E (-) induced a dose-response reduction in sAPPβ in CHO-7W cells at concentrations of 5 nM and higher (**Fig. 6B**), and in Aβ1-42 at concentrations of 25 nM and higher (**Fig. 6C**).

**Figure 6.**
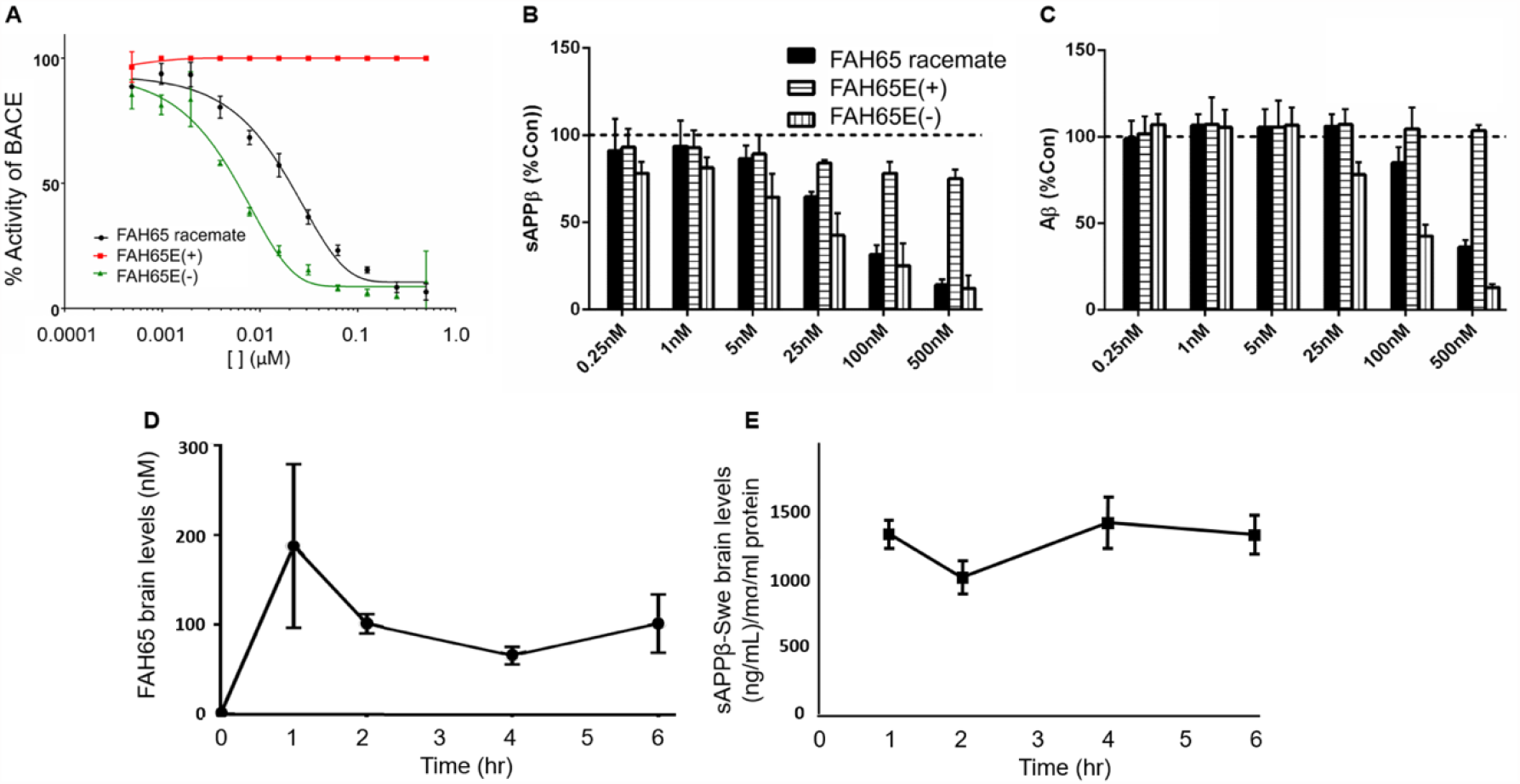
Inhibitory activity and brain permeability of FAH65E(-). Shown are (A) dose-response curves for FAH65 racemate and the FAH65E(+) and (-) enantiomers in the P5-P5’ assay and (B) sAPPβ and (C) Aβ1-42 in CHO-7W cells after treatment with increasing concentrations of FAH65 racemate and enantiomers. Legend in B applies to C. Data graphed as the mean and SEM. (D) Brain levels of FAH65E(-) and (E) sAPP*β* levels in brain from ApoE4TR-5XFAD mice after oral delivery of 30 mg/Kg FAH65E(-). Note: PK study design did not include 0 timepoint untreated mice. N = 3 mice per time point.

In ADME-T testing (see **Supplementary Methods** and **Supplementary Table S2)**, FAH65E(-) displayed favorable physiochemical properties to be an orally bioavailable therapeutic in vivo, including kinetic solubility at 79 μM, potential for blood-brain barrier (BBB) permeability at 1.01 Pm in PAMPA, and plasma stability at t_1/2_ > 180 min. It has a moderately short half-life in liver microsomes (49 min) and bound to HSA at relatively high levels (F_u_ 0.49%). The F_u_ was 5.4% for brain tissue.

In vivo, FAH65E(-) was observed to be orally brain bioavailable, reaching a maximum brain concentration (C_max_) of 188 nM at 1 hour after oral administration at 30 mg/Kg to ApoE4-5XFAD mice (**Fig. 6D**). In the same mice, sAPPβ decreased 2 hours after the oral administration (**Fig. 6E**) showing a correlation between C_max_ and pharmacodynamics effects after administration of FAH65E(-).

## Discussion

We describe herein our discovery of APP-selective BACE inhibitor (ASBI) lead candidates FAH65 and its enantiomer FAH65E(-) through screening, and optimization by exploratory medicinal chemistry, and analog synthesis. The lead candidate FAH65 is highly selective for BACE1 as the target enzyme and APP as the substrate. *In vitro* studies reveal the selectivity and potency of FAH65, and *in vivo* studies in AD model mice provide evidence that FAH65 decreases APP cleavage product sAPPβ in brain tissue and increases trophic, neurite-supporting cleavage product sAPPα, along with significant improvements in memory in the NOR/NLR testing paradigms with oral treatment at 30 mkd for 26 days.

In the pilot study in rats, FAH65 displayed good brain penetrance in brain and target engagement, lowering sAPPβ and Aβ1-40/42, with a concomitant increase in sAPPα, in both CSF and brain. Overall, the effects of FAH65 in the rat study were greater than the clinically-tested BACE inhibitor verubecestat. The potential advantage of FAH65, which has BACE inhibition potency similar to that of veru, is its high selectivity for inhibiting BACE cleavage of APP relative to other substrates such as NRG1 and PSGL1.

FAH65 has good drug-like properties, including stability in human plasma, microsomal stability, and kinetic water solubility. It also has a good hERG profile with an EC50 > 10 μM. The F_u_ of 5.4% for mouse brain tissue protein binding suggests the unbound brain concentration of FAH65-E(-) is 9.6 nM, which is less than the predetermined in-vitro efficacious dose (EC50 = 25 nM). Since the oral administration of FAH65E(-) at a dose of 30 mkd is expected to inhibit BACE1 less than 50% in vivo in the brain. We note ASBI therapy with FAH65 herein may only result in partial, not complete, inhibition of BACE1 in the brain. Such a partial BACE inhibition therapeutic approach is supported by the previous finding of McConlogue et al. (29) that indicated, at least in a mouse model of AD, that only a 12% decrease in Aβ levels results in dramatic reduction in plaque pathology. The potential of modest BACE inhibition and partial reduction of Aβ production in the brain was also suggested by Satir et al. (30), who found that less than a 50% decrease in Aβ secretion by a BACE inhibitor – a magnitude similar to that in the presence of the protective Icelandic mutation (14) - does not impair synaptic function and thus is not likely to cause the cognitive deterioration observed with some BACE inhibitors that have failed in the clinic.

Despite reported target engagement, BACE inhibitors that have been studied in the clinic have not elicited improvements in cognitive performance, and in some instances have been associated with worsened cognitive performance (31). The failure of BACE inhibitors in the clinic to-date have been attributed to off-target effects due to inhibition of non-APP BACE substrates, a limitation that an ASBI specifically addresses and is likely to overcome.

If successfully developed, a small molecule ASBI therapeutic with good oral bioavailability and brain penetrance may also likely have advantages over the currently approved amyloid-targeted therapeutic aducanumab and a similar monoclonal antibody-based therapy such as lecanemab (15, 16) that only provided a modest decrease in decline of memory, in ease of delivery – oral dosing vs. infusion – and safety risks such as inflammation, ARIA, and risk of bleeding in the brain.

In our next steps toward preclinical development of this ASBI lead candidate, the active enantiomer of FAH65E(-), will undergo in vivo testing in murine models to assess effects on BACE activity/substrate cleavage, including off-target effects, in brain as well as other IND-enabling studies.

An ASBI that reaches the clinical testing stage may be most beneficial for patients before the onset of symptoms. For example, in persons who possess ApoE4 alleles or who are otherwise at risk for increase beta processing of APP. Further, an ASBI may also provide the possibility of combination with FDA-approved mAb-based therapy with the potential to increase efficacy of both therapeutics by decreasing Aβ production while clearing existing amyloid; it may also allow a decrease in the dose of the mAb and a reduction in observed clinical side effects and could serve as a maintenance therapy to maintain normal levels after amyloid is cleared from the brain.

## Author contributions

JC performed in vitro assays and analyzed data; BJ participated in developing medical chemistry and analog design, OD performed the screening assay and in vitro studies, WC and JL performed pharmacokinetic and ADME analyses, OG assisted with in vivo studies, KP, MJ, and CE performed assays, CPL performed the APP binding studies, PS performed in vivo studies, analyzed and graphed data and wrote the manuscript, DB and VJ originated the concept of ASBI identification and designed the screening assay, medicinal chemistry strategy and analog generation, in vitro and in vivo studies, analyzed data and edited the manuscript.

## Acknowledgements

We thank Dr. Clare Peters-Libeu, who performed the APP binding studies. This research was supported by funding from the Easton Center for Alzheimer ’s Disease Research and Care at the Department of Neurology at UCLA and a sponsored research agreement to the Drug Discovery Lab at UCLA by NantNeuro LLC. The support by the UCLA Technology Development group (TDG) leading to the issued claims from this research in UCH-14160.

## Methods

### P5-P5’ assay

For the P5-P5’ BACE inhibition assay, 4 μL of assay buffer was added to each well, followed by 2 μL of BACE-1 diluted in assay buffer to 7.5 ng/μL. Then, 2 μL of inhibitor, at concentrations along an 11 pt. two-fold dilution series starting at 10 μM, or 0.5 μM, were added to appropriated wells and incubated for 30 minutes at room temperature. For all dilutions of inhibitor, drug stock solutions of 10 mM in DMSO were diluted to 50 μM in water; subsequent dilutions were in 0.5% DMSO. Afterwards, 2 μL of fluorogenic P5-P5’ BACE-1 substrate, diluted to 50 μM in assay buffer, were added to each well and the signal generated was read every 30 minutes at 25°C for 2hr.

### SEAP-NRG assay

A cDNA construct encoding a human placental secreted alkaline phosphatase (SEAP)-NRG1 (pAPtag5-NRG1-β1) fusion protein was transfected in human embryonic kidney (HEK293) cells in a 6-well format with or without full-length wild-type BACE1 using Lipofectamine 2000 (Invitrogen) as described previously (31). After transfection, the medium was replaced with DMEM containing the test compound at 10, 5, 1, 0.1, and 0.01 μM and 10% heat-inactivated fetal bovine serum (FBS), and incubated for 24 hours. SEAP activity was measured in the conditioned medium. For alkaline phosphatase activity measurements, 200 μL of reaction solution (0.1 M glycine, pH 10.4, 1 mM MgCl2, 1 mM ZnCl2 containing 1 mg/mL 4-nitrophenyl phosphate disodium salt hexahydrate, Sigma) were added to 20 μL of the conditioned medium. The absorbance was read at 405 nm. Statistical analyses were performed using a two-tailed Student’s t-test.

### PSGL1 assay

A cDNA construct encoding a human placental secreted alkaline phosphatase (SEAP)-PSGL1 (pAPtag5-PSGL1-β1) fusion protein was transfected in human embryonic kidney (HEK293) cells in a 6-well format with or without full-length wild-type BACE1 using Lipofectamine 2000 (Invitrogen). After 6-8h transfection, the medium was replaced with DMEM containing the test compound at 10010, 5, 1, 0.1 and 0.01 μM and 10% heat-inactivated fetal bovine serum (FBS), and incubated for 24 hours. SEAP activity was measured in the conditioned medium. For alkaline phosphatase activity measurements, 200 μL of reaction solution (0.1 M glycine, pH 10.4, 1 mM MgCl2, 1 mM ZnCl2 containing 1 mg/mL 4-nitrophenyl phosphate disodium salt hexahydrate, Sigma) were added to 20 μL of the conditioned medium. The absorbance was read at 405 nm at 60 min.

### Cathepsin D assay

The effects of candidate ASBIs on cathepsin D (CatD) activity was assessed using the R&D Systems CatD activity assay kit (Catalog #: 1014-AS and ES001) accordingly to the manufacturer’s instructions. For the assay, 4 μL of assay buffer was added to each well, followed by 2 μL of CatD diluted in assay buffer to 5 ng/uL. Then, 2 μL of inhibitor, at concentrations along an 11 pt. two-fold dilution series starting at 50 μM, were added to appropriated wells and incubated for 30 minutes at room temperature. For all dilutions of inhibitor, drug stock solutions of 10 mM in DMSO were diluted to 50 μM in water; subsequent dilutions were in 2.5% DMSO. Afterwards, 2 μL of fluorogenic CatD substrate, diluted to 50 μM in assay buffer, were added to each well, after which the signal generated was read every 30 minutes at 25°C for 1hr.

### BACE2 assay

The BACE2 assay is described in the **Supplementary Materials**.

### Plasmids

The pAPtag5-NRG1-β1 construct was kindly provided by Dr. Carl Blobel (33). The BACE1 construct was a gift from Dr. Michael Willem and Dr. Christian Haass (32).

### Medicinal chemistry and testing of analogs

Our exploratory medicinal chemistry efforts focused on the phenytoin hit to see if we could generate analogs with enhanced potency. The medicinal chemistry efforts (planned to be published elsewhere) involved conversion of the hydantoin scaffold to an aminohydantoin that led to our potent ASBI FAH65. Analogs were assessed in the MBP-APPC125, p5-p5’, and substrate selectivity assays described below.

### Synthesis

The methods for FAH analog synthesis are described in the **Supplementary Materials**.

### CHO-7W and -7W^Swe^ cell culture

The Chinese hamster ovary (CHO) cell line overexpressing human AβPP (7W) stably transfected with human APP or overexpressing APP with the Swedish (Swe) K595N/M596L mutation were incubated with FAH compounds diluted to 1 µM overnight. Culture Media was collected and frozen. Media was assayed using an AlphaLISA assay for Aβ1-42 (Perkin Elmer catalog # AL276C), sAPPα (R&D Systems catalog # AF1168 conjugated with Perkin Elmer acceptor beads catalog # 6772001 + 2B3 antibody from IBL catalog # 11088 biotinylated), and sAPPβ (Perkin Elmer AL276-acceptor + IBL catalog # 18957 biotinylated). For the assay, 2 µL of diluted or undiluted sample was added to a 384 well plate, followed by 2 µL of the respective antibody mixture. This was incubated for 1hr. Afterwards, 2 µL of donor beads was added to each well, followed by a 30-minute incubation in the dark, after which the signal was read.

### Pharmacokinetics (PK)/Pharmacodynamics (PD) in mice and rats

All in vivo experiments described were carried out in strict accordance with good animal practice according to NIH recommendations. All procedures for animal use were approved by the Animal Research Committee (ARC) at UCLA and under an approved ARC protocol.

Wildtype or non-transgenic (NTg) C57Bl6 mice or Sprague-Dawley rats were dosed by oral (gavage) or subcutaneous (SC) injection at 10 or 30 mg/kg, and animals were euthanized 1, 2, 6, and 8 hours after administration by ketamine/xylazine over-anesthesia following by transcardial collection of blood for isolation of plasma and saline perfusion. Brain tissue was collected post-mortem for assessment of compound levels and biomarkers. In rats, CSF was also collected.

Analyses of compound levels in PK studies were performed in the UCLA Pasarow Mass Spectrometry Lab (PMSL; Julian Whitelegge, Ph.D., Director). Tissues were homogenized in a bead beater using 5 volumes of ice-cold 80% acetonitrile (1/5; mg of brain/µL of 80% ACN). Plasma analytes were extracted using 4 volumes of ice-cold acetonitrile (1/4; µL of plasma/µL of 100% ACN). Solutions were clarified by centrifugation (16,000 x g, 5 min) and the supernatants were transferred to new tubes and lyophilized. Samples were reconstituted in 100 µL of 50/50/0.1 (Water/Acetonitrile/Formic Acid) prior to analysis via liquid chromatography-tandem mass spectrometry (LC-MS/MS).

An internal standard (IS) was added to every sample to account for compound loss during sample processing. Standards were made in drug naïve plasma and tissue lysates with increasing amounts of analyte (S1, S2: 0 pmol/ S3, S4: 1 pmol/ S5,S6: 10 pmol/ S7,S8: 100 pmol, S9,S10: 1000 pmol). The standard curve was made by plotting the known amount of analyte per standard vs. the ratio of measured chromatographic peak areas corresponding to the analyte over that of the IS (analyte/IS). The trendline equation was then used to calculate the absolute concentrations of each compound in plasma and tissue.

The targeted LC-MS/MS assay was developed using the multiple reaction monitoring (MRM) acquisition method on a 6460 triple quadrupole mass spectrometer (Agilent Technologies) coupled to a 1290 Infinity HPLC system (Agilent Technologies) with a Phenomenex analytical column (Kinetex 1.7 µm C18 100 Å 100 × 2.1 mm). The HPLC method utilized a mixture of solvent A (99.9/1 Water/Formic Acid) and solvent B (99.9/1 Acetonitrile/Formic Acid) and a gradient was use for the elution of the compounds (min/%B: 0/1, 3/1, 19/99, 20/1, 30/1). Two fragment ions were monitored at specific LC retention times to ensure specificity and accurate quantification in the complex biological samples. The normalized chromatographic peak areas were determined by taking the ratio of measured chromatographic peak areas corresponding to the compound over that of the internal standard (Analyte/IS).

The T_max_, C_max_, brain-to-plasma ratios and brain levels were then calculated using PK Solutions software (SummitPK).

### Efficacy testing in B254 mice

B254 mice are an AD model comprising expression of human APP with Swedish and Indiana mutations along with a D664A substitution, as described in Galvan et al. (34). Male and female 4 to 5 month-old B254 mice were administered FAH65 via pipette feeding by the method of Atcha et al. (35) at a dose of 30 mg/kg BID (60 mg/kg/day). Mice were weighed before the first dose for the calculation of volume to be administered. The formulation used comprised FAH65-HCl as a 40 mg/mL solution in water plus 30 mg/mL in strawberry syrup.

In the pilot study, mice were treated for 10 days, and in the follow-up study, for 26 days. One week before the end of the study, mice in the pilot study underwent Novel Object Recognition (NOR) memory testing, and mice in the follow-up study underwent both NOR and Novel Location Recognition (NLR) testing as described elsewhere (28, 36). Briefly, mice are introduced to two identical, parallel and evenly objects in an opaque 30 × 40 cm arena (box without a lid) with navigation marks on the wall, for 10 min. For Novel Location Recognition (NLR), after a 2.5-hour interval, the mouse is returned to the arena wherein one identical object has been moved diagonally. The time spent and number of discreet interactions with the objects in the original location and in the new location over 10 min is recorded. The mouse is then removed from the arena, and after an additional 2.5 hour interval, for the Novel Object Recognition (NOR) phase, is returned to the arena wherein the objects are in the original position, but one is a new, different object. The time spent and number of discreet interactions with the objects in the original location and in the new location over 10 min is recorded. The novelty preference was calculated by dividing total interactions (or time) by interactions (or time) with the object in the novel location or the novel object.

### Biomarker analyses

Homogenization and BCA: For in vivo studies, Individual tissues were weighed and sonicated on ice with AlphaLISA Lysis Buffer complemented with HALT (Fisher catalog # 78446) to give a 10% W/V sonicate. Total protein concentration was assessed by BCA assay (Fisher catalog # 23227) following manufacturer recommendations

### sAPPβ

For the ELISA assay to assess sAPPβ levels in brain an IBL ELISA (kit #27733) specific for sAPPβ with the Swedish (K595N/M596L) mutation (12) that is present in the APP transgene of the mouse model used following the kit instructions. Briefly, 499 µL of EIA buffer was added to individual tubes for each sample, 1 µL of sample added, vortexed and centrifuged. To each assay plate well, 100 µL of sample was added and the covered plate incubated overnight at 4°C. The plate(s) was washed 4 times with 350 µL/well wash buffer, then 100 µL/well of labeled antibody added followed by incubation for 30 minutes at 4°C. After five washes, 100 µL chromogen solution was added to each well and incubated for 30 min. at RT in the dark. Stop solution (100 µL/well) was then added and the plate read at 450 nm.

### βCTF

The BACE cleavage product βCTF was detected by AlphaLISA (Perkin Elmer) Amyloid Beta kit (Cat # AL275C) modified by replacing the anti-Aβ acceptor beads (Cat # AL275AC) specific for the C-terminus of Aβ40 with anti-Aβ acceptor beads from the AL202 kit (Cat #AL202AC) having the 4G8 antibody. For the assay, 2 µL of diluted sample was added to a 384 well plate, followed by 2 µL of the antibody mixture. This was incubated for 1hr. Afterwards, 2 µL of donor beads was added to each well, followed by a 30-minute incubation in the dark, after which the signal was read.

### sAPPα

sAPPα was assessed by AlphaLISA comprising antibody AF1168 (R&D Systems catalog # AF1168) conjugated with Perkin Elmer acceptor beads (catalog # 6772001) and 2B3 antibody from IBL catalog # 11088 biotinylated and streptavidin donor beads. For the assay, 2 µL of diluted or undiluted sample was added to a 384 well plate, followed by 2 µL of the respective antibody mixture. This was incubated for 1hr. Afterwards, 2 µL of donor beads was added to each well, followed by a 30-minute incubation in the dark, after which the signal was read.

### Aβ1-42

For assessment of Aβ 1-42 in brain, tissues were weighed and sonicated in freshly prepared 5 M Guanidine (Gdn)-HCl in 50 mM Tris–NCl pH 8.0 at 20% weight/volume. Samples were rotated for 2–3 h at room temperature after sonication and typically frozen stored before use. An Invitrogen ELISA kit for 1-42 was used according to the manufacturer’s instructions.

### Total tau and p-tau

Total tau and p-tau were assessed by Perkin-Elmer AlphaLISA kits AL271 and AL3136, respectively. The assays were run as described above for the sAPPa AlphaLISA.

### FAH65 enantiomer separation

Methods for FAH65 enantiomer separation are described in **Supplementary Methods**.

### ADME-T

Methods for determination of kinetic solubility, plasma stability, liver microsome stability, parallel artificial membrane permeability assay (PAMPA), plasma protein (human serum albumin, HSA) binding, and brain tissue binding, as well as hERG analysis, are described in **Supplementary Methods**.

