## Supplementary Material, Table S1-S3 and Fig S1-S42 for "Discovery of an APP-Selective BACE1 Inhibitor for Alzheimer’s Disease"

**FAH1 and FAH3 Dose-response**……………………………………………… …………….………..2

**Figure S1.** *IC50 curve for FAH1, 3, 17 and 65*…………………………………………………………….2

**Table S1.** *Selectivity of BACE inhibitors*.........................................................................................2

**BACE2 assay**……………………………………………………………………………..………………2

**Figure S2.** *FAH65 in BACE2 assay*……………………..……………………………………………..……3

**ADME-T assays**………………………………………………………………………………..…………..…3

**Table S2.** *Characteristics of FAH65 racemate & enantiomers.*……………………………….…..………5

**Table S3.** *FAH binding to APP*…………………………..…………………………………….…..………5

**hERG assay of FAH65**………………………………………………………………………………...……..5

**APP- binding to FAHs**…………………………………………………………………………………….5

**FAH65 vs. verubecestat in rats**…………………………………………………………………...……5

**Figure S3**. FAH65 vs. veru in rats……….………………………………..………………………...………6

**SFC separation of FAH65 enantiomers**……………...….……………………………………………..……7

**Optical rotation of FAH65 enantiomer**s…………………………………………………………………..8

**Synthesis of analogs**

*FAH1*…...……………………..……………………………………………………………………………..9
*FAH3*……………………………………………………………………………………………………..….13
*FAH17*...…………………………………………..…………………………………………………...……18
*FAH65*.........................................................................................................................................26
*FAH65E(+)*.....................................................................................................................................31
*FAH65E(-)*......................................................................................................................................33

#### FAH1 and FAH3

The IC50s in the P5-P5’ assay for FAH1 and 3 were 5-10 µM and 1-4 µM, respectively (**Figure S1A**). In CHO-7W cell culture, both decreased Aβ42, FAH1 at 5 µM and FAH3 at 1 µM. Evaluation of FAH3 in SEAP-NRG assay shows that it had selectivity for inhibiting BACE-APP cleavage, where 50% inhibition was seen at ~1uM, while no inhibition observed for BACE-NRG cleavage at 1uM. Both showed brain permeability after subcutaneous (SubQ) injection at 10mg/Kg (**Figure S1B &C**).

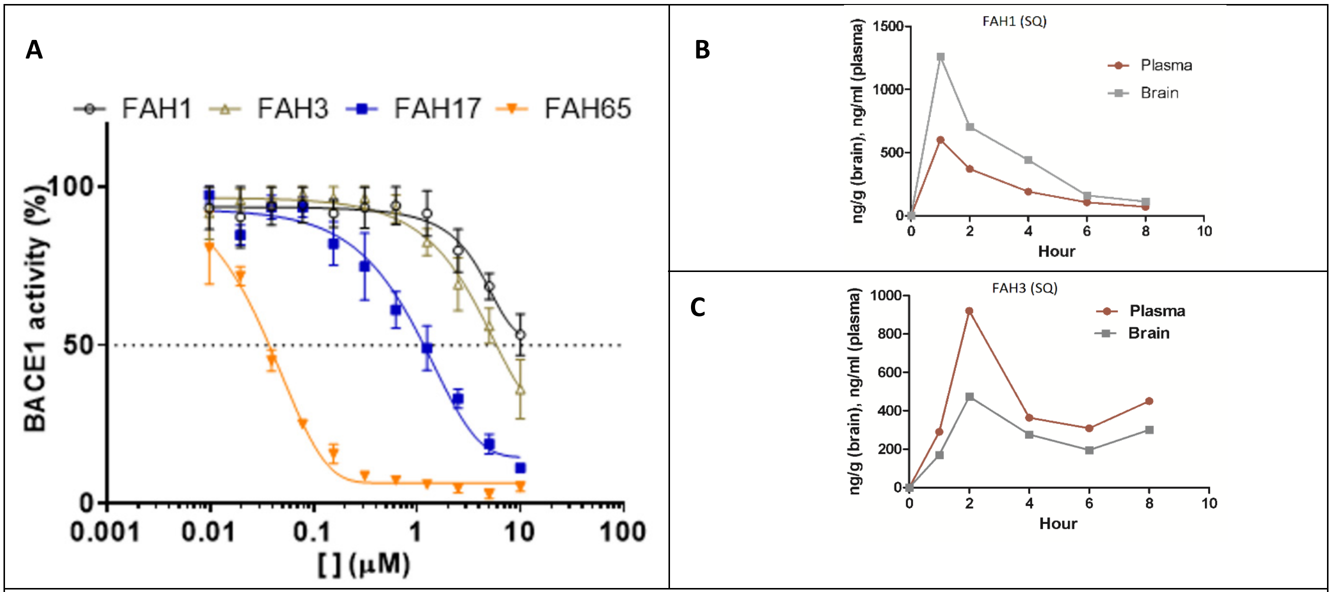


**Figure S1A.** *IC50 curves for FAH1, 3, 17 and 65 in the P5-P5’ assay.* **Figure S1B &C.** *PK analysis to determine drug levels in brain and plasma after 10mg/Kg (SubQ) injection of FAH1 and FAH3, n=1 number of mice per time point.*

#### Selectivity of BACE inhibitors

**Table S1. Selectivity of BACE Inhibitors**

| P5-P5' (IC50, µM) | | | NRG1 (IC50, µM) | | | IC50 NRG1/IC50 APP | | |
|---|---|---|---|---|---|---|---|---|
| FAH65 | Lana | Veru | FAH65 | Lana | Veru | FAH65 | Lana | Veru |
| 0.007 | 0.004 | 0.002 | 2 | < 0.01 | <0.01 | > 200 | <2.5 | <2.5 |

Lana - lanabecestat; Veru - verubecestat

#### BACE2 assay

The effects of candidate ASBIs on BACE2 activity was assessed using the R&D Systemes BACE2 activity assay kit (Catalog #: 4097-AS and ES010) formatted from to the manufacturers. For the assay, 4 µL of assay buffer was added to each well, followed by 2 µL of BACE2 diluted in assay buffer to 10 ng/µL. Then, 2 µL of inhibitor, at the appropriate concentrations were added to the wells and incubated for 30 minutes at room temperature. For all dilutions of inhibitor, drug stock solutions of 10 mM in DMSO were diluted to 50 µM in water; subsequent dilutions were in 0.5% DMSO. Afterwards, 2 µL of fluorogenic BACE2 substrate, diluted to 50 µM in assay buffer, were added to each well, after which the signal generated was read every 30 minutes at 25°C for 2 hr.

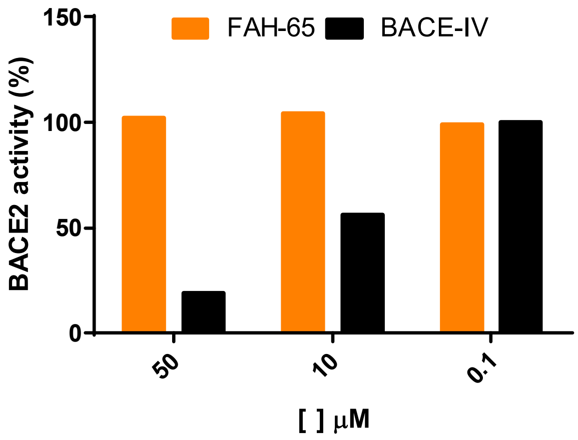


**Figure S2.** *BACE2 activity.* FAH65 shows no inhibition of BACE2 activity.

### ADME-T assays

*Kinetic solubility assay*

FAH65E at 10 mM in 100% DMSO was diluted in PBS pH 7.4 to 100, 10, 1, and 0.1 µM; plus a vehicle-only control. The solutions were incubated at 37°C for 90 min and centrifuged at 16K x g for 5 min. An aliquot of each supernatant was analyzed by LC-MS/MS. A standard curve was generated by plotting the known amount of analyte per standard in DMSO vs. chromatographic peak area.

Kinetic solubility (mM) was calculated using the trendline equation with maximum chromatographic peak area observed in the aqueous sample.

*Liver microsome stability assay*

A 1 µL sample of 1 mM FAH65E in 100% DMSO was added to 1 mL of 0.5 mg/mL human liver microsomes (Thermo Fisher Scientific, Cat# HMMPL) in PBS pH 7.4, 2 mM NADPH, and 2 mM MgCl2; and incubated at 37°C for 120 min. Aliquots (50 µL) of the microsome solution were taken at timepoints of 0, 5, 10, 15, 30, 60, 90, and 120 min, and added to a 200 µL 100% acetonitrile reaction quenching solution containing an internal standard. Solutions were clarified by centrifugation (16K x g, 5 min) and supernatants transferred to new tubes and lyophilized. Samples were reconstituted in 100 µL of 50/50/0.1 water/acetonitrile/formic acid prior to analysis via LC-MS/MS.

Normalized chromatographic peak areas were plotted at each time point and the half-life ($T_{1/2}$) of compound in liver microsomes was determined by using the trendline equation to calculate the time at which compound abundance was 50% of that at timepoint 0 (T0).

*Plasma stability assay*

One µL FAH65E (1 mM in 100% DMSO) was added to human peripheral blood plasma (Cat#70039, Stemcell Technologies Inc) diluted 2-fold in PBS (1000 µL, pH 7.4) and incubated at 37°C for 180 min. Aliquots (50 µL) of the plasma solution were taken at 0, 15, 30, 60, 90, 120, 150, and 180 min, and added to a 200 µL 100% acetonitrile reaction quenching solution containing an internal standard. Solutions were clarified by centrifugation (16K x g, 5 min) and supernatants were transferred to new tubes and lyophilized. Samples were reconstituted in 100 µL of 50/50/0.1 water/acetonitrile/formic acid prior to analysis via LC-MS/MS).

Normalized chromatographic peak areas were plotted at each time point and the $T_{1/2}$ of FAH65E in plasma was determined by using the trendline equation to calculate the time at which compound abundance was 50% of that at timepoint 0 (T0).

*Human serum albumin (HSA) binding assay*

The liquid chromatography-ultraviolet/visible spectroscopy (LC-UV/Vis) assay was performed on a 1290 Infinity HPLC system (Agilent Technologies) with an HPLC column containing immobilized HSA (CHIRALPAK® CAN, Chiral Technologies, Cat#H15L-109, 5 µm 50 x 4 mm). The HPLC method utilized an isocratic gradient containing 100% 6.7 mM phosphate buffer saline (pH 7.4). The retention time of the compound (tr) and injection peak were recorded.

The percent protein binding (P) was calculated using the following equations:

$K' = (t_r - t_m) / t_m$; $P = 100 \times [K' / (K' + 1)]$

*Parallel artificial membrane permeability assay (PAMPA)*

A Regis A liquid chromatography-ultraviolet/visible spectroscopy (LC-UV/Vis) assay was performed on a 1290 Infinity HPLC system (Agilent Technologies) with an HPLC column containing immobilized phosphatidylcholine (IAM.PC.DD, Regis Technologies, Cat#774011, 5 µm 300 Å 100 x 4.6 mm). The HPLC buffer comprised 6.7 mM PBS (pH 7.4; solvent A) and acetonitrile (solvent B): a gradient was used for the elution of the compounds (min/%B: 0/20, 20/60, 21/20, 30/20). The retention time of the compound (tr) and void volume time of the column (t0) were recorded.

Membrane permeability (Pm) was calculated using the following equations as described by Yoon et al (DOI: 10.1177/1087057105281656):

$K_{IAM} = (t_r - t_0) / t_0$; $Pm = (K_{IAM} / MW^4) \times 10^{10}$

Compounds with a Pm >0.85 are predicted to be BBB permeable (CNS+) at pH 7.4.

*Brain Tissue Binding Assay*

Brain tissue was homogenized in PBS (pH 7.4) (1: 3 weight(mg)/volume(µL)) and the protein concentration determined using the Micro BCA™ Protein Assay Kit (Thermo Fisher Scientific, Cat#23235). Brain homogenate was diluted to 20 mg/mL in PBS (pH 7.4) and added to Slide-A-Lyzer™ MINI Dialysis Devices, 10K MWCO dialysis cups (Thermo Fisher Scientific, Cat#PI88401) in a 48-well plate containing PBS (500 µL; pH 7.4). 1 µL of 1 mM compound was added to the brain homogenate (final concentration: 2 µM compound, 0.5% DMSO) and incubated on a rocker for 4.5 hours at 37 °C. 50 µL of brain homogenate (within the dialysis cup) and PBS (within the 48-well plate) were transferred to new microcentrifuge tubes containing 200 µL of quenching reagent (100% acetonitrile) containing internal standard. Solutions were clarified by centrifugation (16,000 x g, 5 min) and the supernatants were transferred to new tubes and lyophilized. Samples were reconstituted in 100 µL of 50/50/0.1 (Water/Acetonitrile/Formic Acid) prior to analysis via LC-MS/MS.

The % of the unbound drug was calculated using the following equation:

$f_{ubound}$ = [normalized peak area in buffer side / normalized peak area in brain side] x 100

| **Table S2. FAH65 Racemate and Enantiomer Characteristics** | | | |
|---|---|---|---|
| Compound | FAH65 Racemate | FAH65E(-) | FAH65E(+) |
| Molecular weight | 423.42 | 423.42 | 423.42 |
| Kinetic solubility (μM) | 88 | 79 | 87 |
| Microsomal stability $T_{1/2}$ (min) | 38 | 24 | 24 |
| Plasma stability $R_{1/2}$ (min) | >180 | >180 | >180 |
| PAMPA Predicted BBB permeability, Pm | 1.09 | 1.01 | 1.08 |
| Oral Brain Levels; $C_{max}$ (nM)* | 188 | 188 | NA |
| Brain:plasma ratio* | 1 : 1.76 | 1 : 1.8 | NA |
| Plasma binding, $f_{unbound}$ (%) | 0.44 | 0.49 | 0.8 |
| Brain tissue binding, $f_{unbound}$ (%) | 3.1 | 5.4 | 6.2 |
| Unbound, $C_{max}$ (nM) | 9.6 | 9.6 | NA |
| In vitro IC50 (nM) | 10-25 | 5 | >10,000 |
| BBB: blood-brain barrier; *PK not performed for E(+) | | | |

*Desired ADME property values: Kinetic Solubility > 5 uM; Microsomal Stability t1/2 > 1 hrs; Plasma Stability t1/2 > 3 hrs; HSA Binding < 95%; PAMPA Pm > 0.85 = CNS+; Brain Tissue Binding fu, bound > 5%.

#### hERG testing of FAH65

The hERG study was conducted at Eurofins (CEREP PanLabs). FAH65 was tested at 1.0E-05 M. Compound binding was calculated as a % inhibition of the binding of a radioactively labeled ligand specific for each target. While results showing an inhibition or stimulation higher than 50% are considered to represent significant effects of the test compounds, such effects were not observed at any of the receptors studied. The range of inhibition for FAH65 was 38.7 to 41% of the potassium channel hERG (human) –[3H] dofitilide assay.

#### APP binding experiments

For these pilot experiments eAPP as described in Descamps et al (26) is crosslinked to a the CM5 Biacore chip. FAH3 at various concentration was used in the flow through the chip, and the plasmon resonance signal was determined using a Biacore T100. The raw data uncorrected for contribution of running buffer is fit to equilibrium binding data and corrected for the contribution of the running buffer to obtain binding constant $K_D$ for FAH3, in contrast for BACE-IV no binding was observed even at 50uM. For FAH17 and FAH 65 APP binding was determined using SPR to determine competition with BSA that non-specifically binds to the eAPP-Biacore chip. The data is shown in **Supplementary Table S3.**

| **FAH3** | **FAH17*** | **FAH65*** | **BACE-IV** |
|---|---|---|---|
| $K_D$ ~ 0.3uM | $K_D$ ~3.3uM (Relative to BSA) | Binding to eAPP-Biacore chip observed relative to BSA but no $K_D$ was calculated | $K_D$ > 50 uM |
| **Table S3:** FAH binding to APP ; * Binding determined in competition with BSA. | | | |

**FAH65 vs. verubecestat in rats**

FAH65 was compared with known BACE inhibitor verubecestat (veru; Merck) in rats in a pharmacokinetic (PK)/pharmacodynamic (PD) study to assess relative brain permeability and target engagement. Adult male and female Sprague-Dawley rats (≥ 200 g) were dosed at 30 mg/kg/day with either FAH65 or veru in 20% hydroxypropyl-β-cyclodextrin by oral gavage. Animals were euthanized at 1, 2, 3, 6, 12 and 24 hours (n = 1/time point; 2 untreated controls) post-dose by pentobarbitol over-anesthesia followed by collection of cerebrospinal fluid (CSF), transcardial blood collection for the isolation of plasma, perfusion with cold saline, and collection of brain tissue (medial cortex) for biochemical analyses.

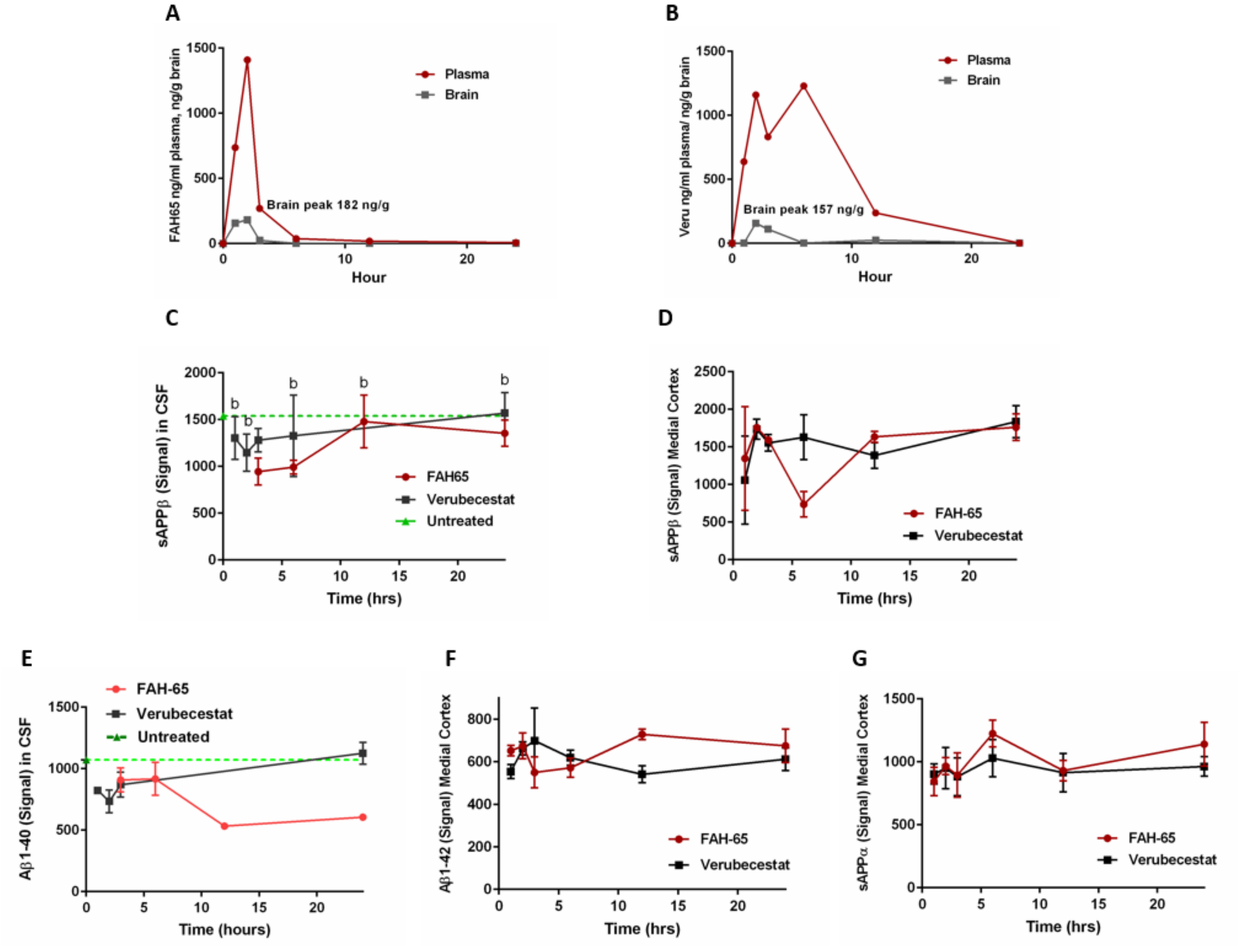


**Figure S3**. *FAH65 and verubecestat (veru) in rats*. (A) FAH65 and (B) veru levels in rat brain tissue (ng/g) and plasma (ng/mL) after oral dosing at 30 mg/kg. sAPPβ in (C) CSF and (D) medial cortex; (E) Aβ1-40 in CSF; (F) Aβ1-42 in medial cortex; and (G) sAPPα levels in medial cortex. N = 3 rats per time point. Blood contamination of some CSF samples prevented accurate assay.

### Separation of enantiomers of FAH65 by Super fluid critical (SFC) chromatography

Enantiomers of FAH65 were separated to generate sufficient amounts for *in vitro and in vivo* studies at Lotus Separations (Princeton, NJ). FAH65 enantiomers were separated by SFC chromatography using a phenomenex column. Separation of 1g of FAH65 yielded 525 mg of the (+)/peak 1 and 465 mg of the (-)/peak 2 enantiomers. In the preparative method a 2 x 25 cm IC was used with 40% isopropanol (0.1% DEA)/$CO_2$ at 100 bar, 50 mL/min, 220 nm with an injection volume of 1 mL, 20 mg/mL ethanol:DCM. For the analytical method, a 25 x 0.46 cm IC was used with 40% isopropanol (DEA)/$CO_2$, at 120 bar3 mL/min, 220, 254 and 280 nm

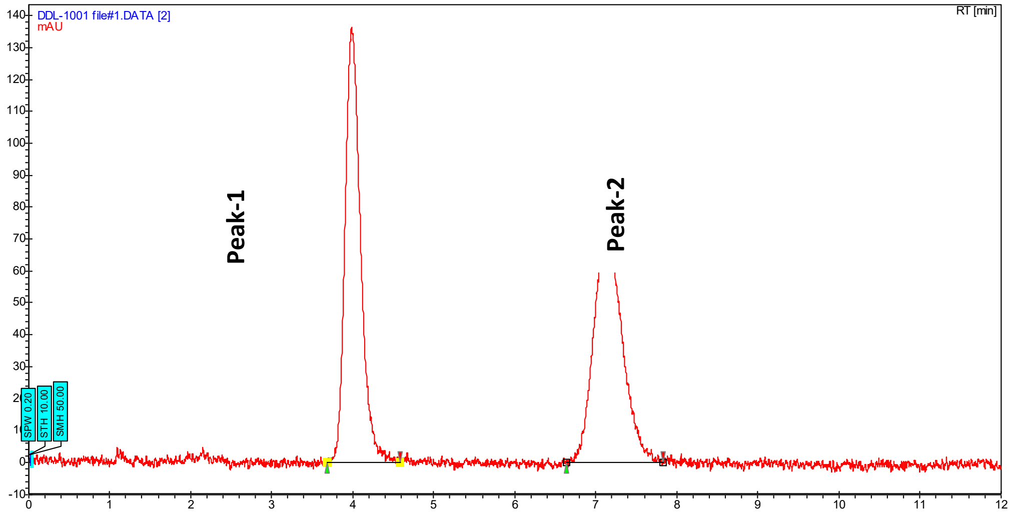


**Figure S4**. *Peaks from 1g starting FAH65.*

**FAH-65 peak-1**

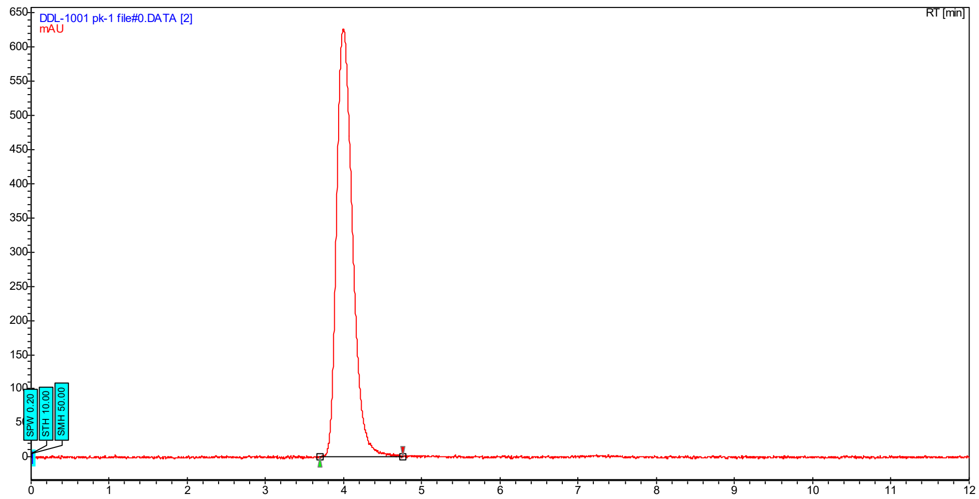


**Figure S5.**

| Index | Time (min) | Area (%) 220 nm |
|---|---|---|
| Peak-1 | 3.99 | 100.00 |
| Total | | 100.00 |

**FAH-65 peak-2**

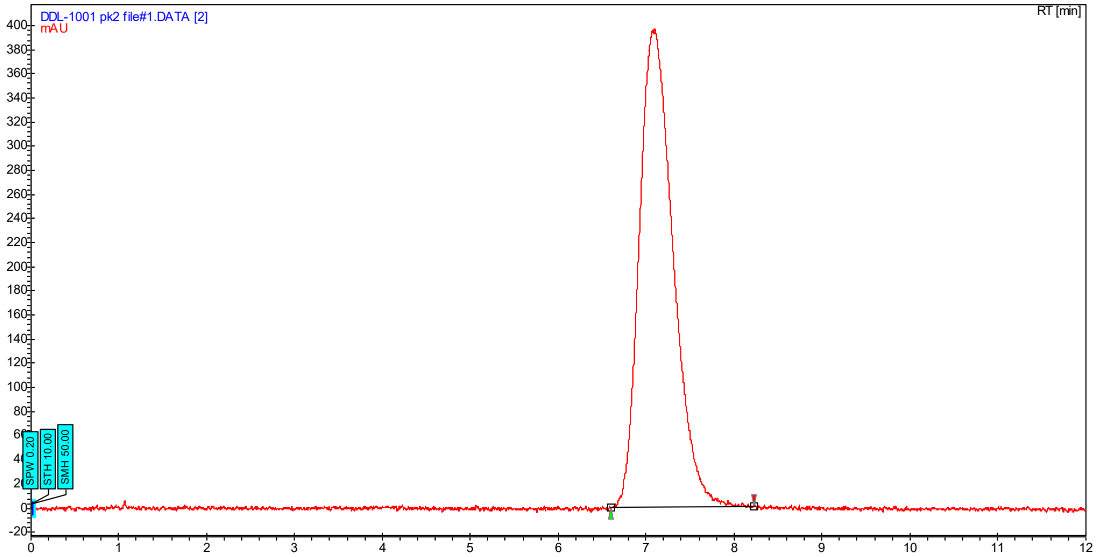


**Figure S6.**

| Index | Time (min) | Area (%) 220 nm |
|---|---|---|
| Peak-2 | 7.09 | 100.00 |
| Total | | 100.00 |

### Optical Rotation

Optical rotations of the FAH65 enantiomers were measured with a Rudolph Autopol III Automatic Polarimeter:

FAH65E(-) (Peak-1): [α]27.0D -180.0° (*c* = 1.00, CH2Cl2).

FAH65E(+) (Peak-2): [α]26.6D +120.0° (*c* = 1.00, CH2Cl2).

### Synthesis of analogs

*FAH-1*

**Step 1:**

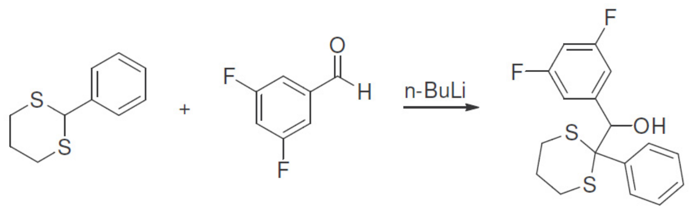


**Figure S7**. *Step 1 of FAH1 synthesis*.

The 2-phenyl-1,3-dithiane (1.79g, 9.12 mmol) was dissolved in 20 mL of dry THF and cooled to 0 degrees. n-BuLi (6.84Ml, 10.94mmol) was added dropwise under nitrogen and the mixture was stirred for 30 min at 0 degrees. A solution of 3,5-difluorobenzaldehyde (1.00mL, 9.12 mmol) in THF (10ml) was added, and the mixture was stirred for 30 minutes, then warmed to ambient temperature over 1 hour and quenched with saturated ammonium chloride solution. The product was extracted into DCM, washed with brine and dried with sodium sulfate. The solvent was removed in vacuo to give (3,5-difluorophenyl)(2-phenyl-1,3-dithian-2-yl)methanol (3.63 g, 8.58 mmol, 94% yield) as a thick yellow oil that solidified on standing. HPLC retention time: 4.496 min, LRMS (M+23) 360.9.

**Step 2:**

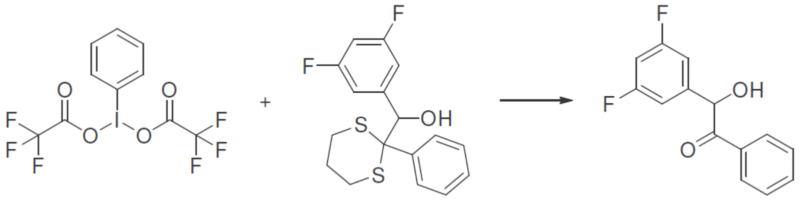

**Figure S8**. *Step 2 of FAH1 synthesis*.

(3,5- Difluorophenyl)(2-phenyl-1,3-dithian-2-yl)methanol (2.90g, 8.58 mmol) was dissolved in 15 mL of acetonitrile and 2.5 mL of water. Bis-(Trifluoroacetoxy)iodobenzene (5.09g, 11.84 mmol) in 10mL of acetonitrile was slowly added at ambient temperature to the vigorously stirred solution. After 1 hour, EtOAC (20mL) was added and the mixture was rinsed with saturated sodium bicarbonate solution followed by brine. The organic fractions were dried, and the solvent was removed in vacuo. The crude product was purified by column chromatography (10-25% EtOAC/hexane) to give 2-(3,5-difluorophenyl)-2-hydroxy-1-phenylethanone (1.29g, 5.20mmol, 60.6 % yield). HPLC retention time: 4.163 min, LRMS (M-1) 247.2.

**Step 3:**

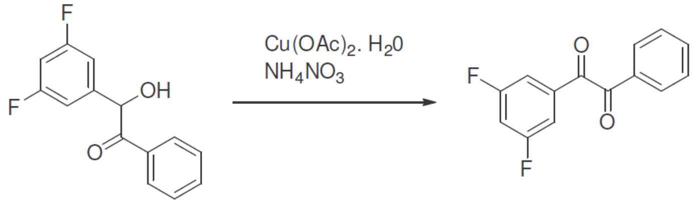


**Figure S9**. *Step 3 of FAH1 synthesis*.

2-(3,5-Difluorophenyl)-2-hydroxy-1-phenylethanone (1.290 g, 5.20 mmol) was dissolved in 80% acetic acid together with diacetoxycopper hydrate (0.052 g, 0.26 mmol) and ammonium nitrate (0.125 g, 1.56 mmol). The mixture was refluxed for 90 minutes and then cooled and the solvent was removed in vacuo. The crude material was purified by column chromatography (5% EtOAcƒhexane) to give 1-(3,5-difluorophenyl)-2-phenylethane-1,2-dione (550 mg, 2.23 mmol 43.0 % yield) as a yellow oil. HPLC retention time: 4.512 min, LRMS (M−1) 245.2.

**Step 4:**

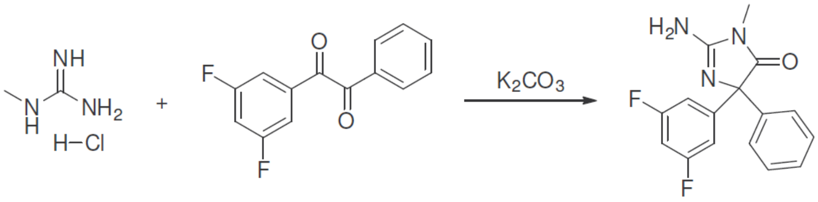


**Figure S10**. *Step 4 of FAH1 synthesis*

To a solution of 1-(3,5-fluorophenyl)-2-phenylethane-1,2-dione (550 mg, 2.23 mmol) in ethanol (20 mL) and water 5 mL) was added 1-methyl guanidine hydrochloride (245 mg, 2.23 mmol) and potassium carbonate (926 mg, 6.70 mmol). The mixture was heated at reflux for 3 hours and then cooled to ambient temperature. The volatiles were removed in vacuo and the residue was taken up in water and extracted into chloroform. The organic fractions were dried with sodium sulfate and the solvent was removed in vacuo. The crude material was purified by preparative TLC (100% EtOAC) to give 2-amino-4-(3,5-fluorophenyl)-1-methyl-4-phenyl-1H-imidazol-5(4H)-one (200mg, 0.66mmol, 29.7%). $^{1}$H NMR (400 MHz, d6-DMSO) δ 7.40-7.06 (m, 8H), 6.76 (s, 2H), 2.95 (s, 3H). retention time: 3.685 min, LCMS (M+1) 302.1.

*FAH1 data analysis*

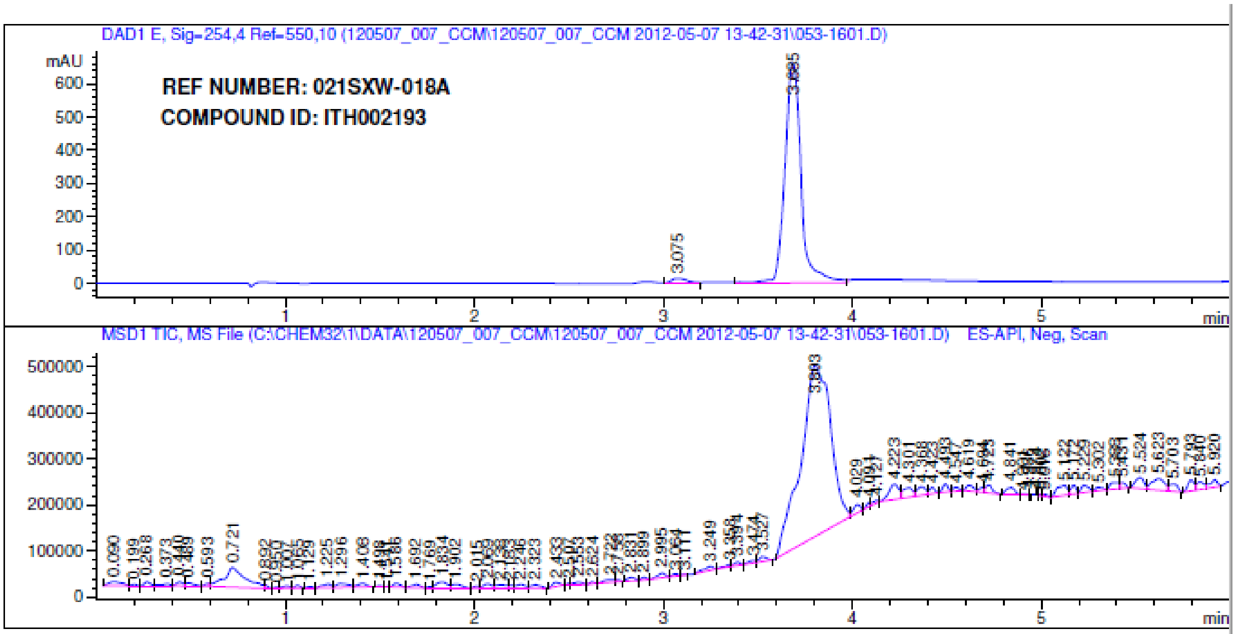


```
Area Percent Report

Sorted By              :      Signal
Multiplier:                    :      1.0000
Dilution:                      :      1.0000
Do not use Multiplier & Dilution Factor with ISTDs


Signal 1: DAD1 E, Sig=254,4 Ref=550,10
```

| Peak # | RetTime [min] | Type | Width [min] | Area [mAU*s] | Height [mAU] | Area % |
|---|---|---|---|---|---|---|
| 1 | 3.075 | VV | 0.0921 | 88.18020 | 14.67076 | 2.1902 |
| 2 | 3.685 | VV | 0.0904 | 3937.94995 | 661.60541 | 97.8098 |

Totals : 4026.13015 676.27616

**Figure S11**. *LC: Chromatogram and Results for FAH1.*

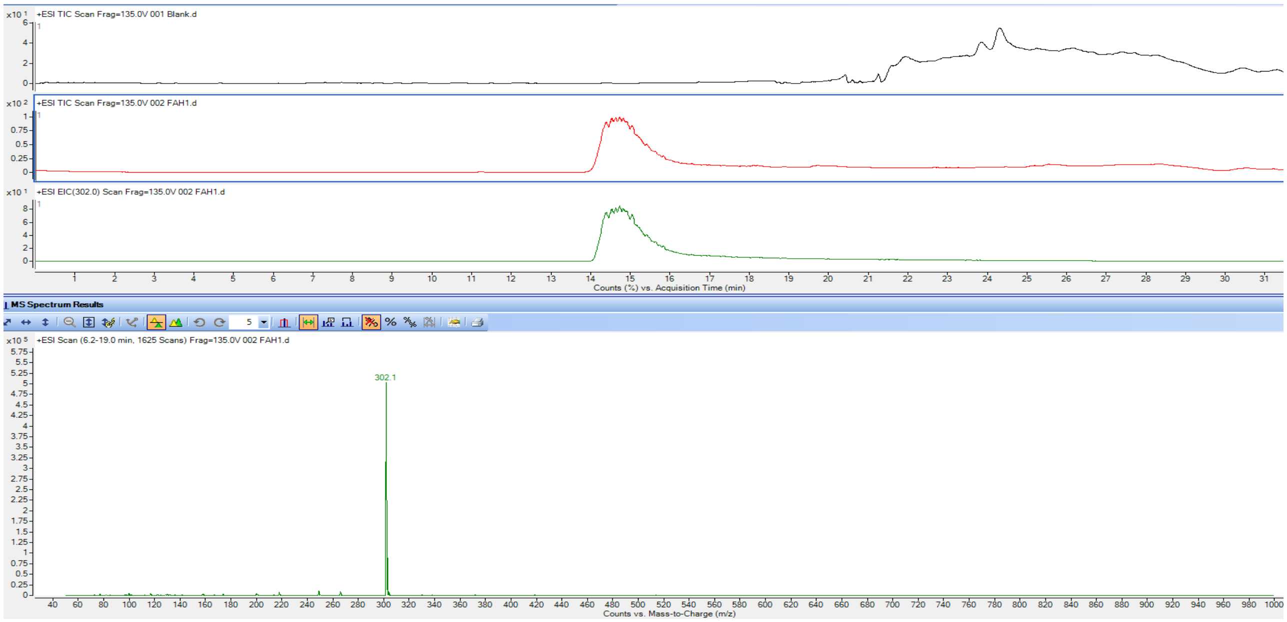


**Figure S12**. *LC-MS for FAH1. Blank TIC in black, FAH1 TIC in red, and FAH1 EIC in green*.

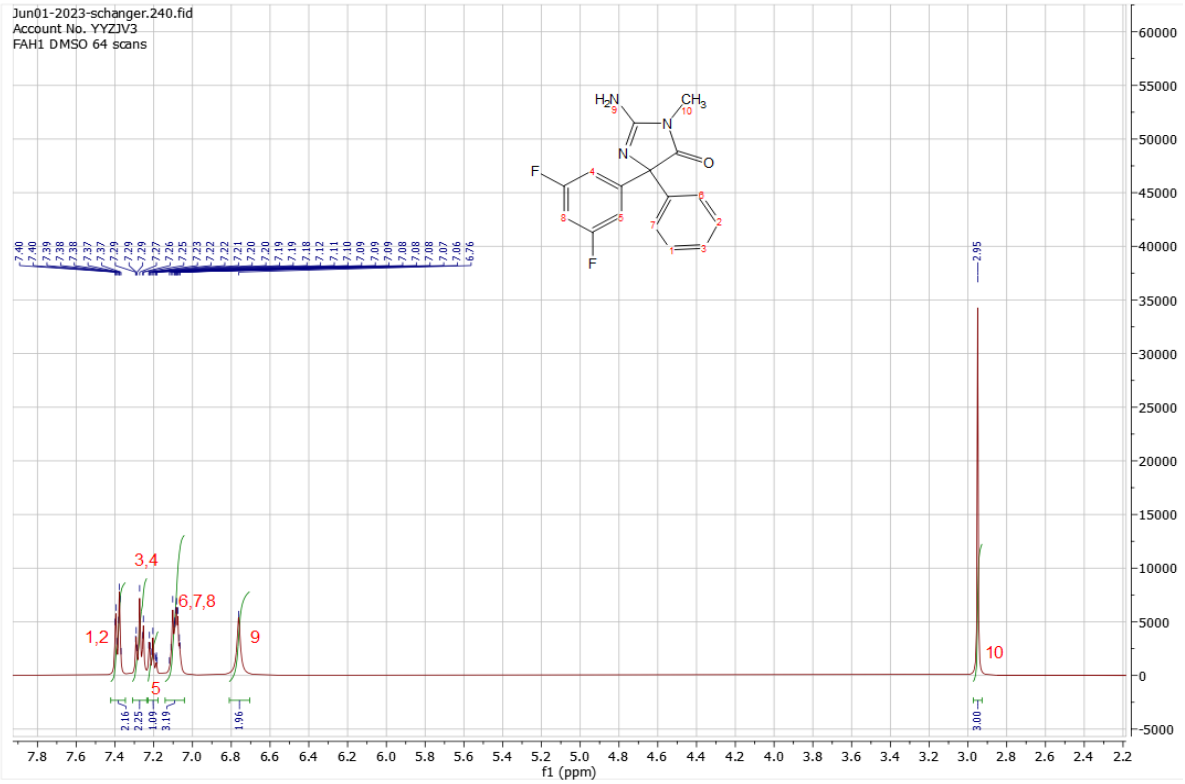


**Figure S13**. *H-NMR for FAH1*

*FAH3*

**Synthesis of 2-(3,5-difluorophenyl)-1,3-dithiane**

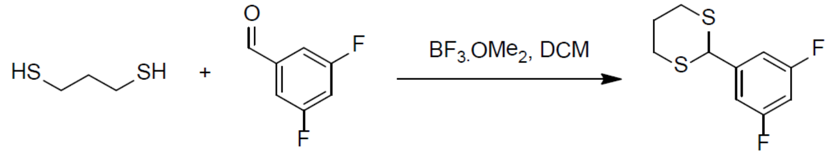


**Figure S14**. *Synthesis of 2-(3,5-difluorophenyl)-1,3-dithiane*.

$BF_3.OMe_2$ (1.40 ml, 15.25 mmol) was added dropwise to a solution of 1,3-propanedithiol (1.81 ml, 17.87 mmol) and 3,5-difluorobenzaldehyde (2.00 ml, 17.87 mmol) in DCM (50 ml) at 0 °C. The reaction was stirred at ambient temperature for 1 hour where TLC (5% EtOAc/hexane) indicated a complete reaction. The reaction mixture was then diluted with additional DCM (50 ml), filtered through Celite, and the Celite pad was washed with additional DCM (3 x 20 ml). The filtrate was washed with saturated NaHCO3 (3 x 50 ml), 10% KOH solution (2 x 50 ml), water (2 x 50 ml) and brine (50 ml) and finally dried over sodium sulfate. The organic extract was filtered and evaporated to afford 2-(3,5-difluorophenyl)-1,3-dithiane (4.12 g, 17.73 mmol, 99%) as a white solid. The proton NMR was consistent with the proposed structure.

**Synthesis of 4-(difluoromethoxy)-3-methylbenzaldehyde**

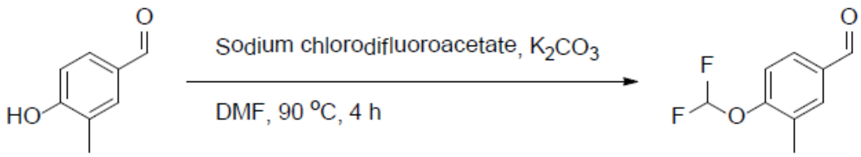


**Figure S15**. *Synthesis of 4-(difluoromethoxy)-3-methylbenzaldehyde*.

A solution of sodium chlorodifluoroacetate (2.60 g, 17.04 mmol) and 4-hydroxy-3-methylbenzaldehyde (1.16 g, 8.52 mmol) in DMF (15 ml) was added over 3 hours to a solution of DMF (15 ml) containing potassium carbonate (1.77 g, 12.78 mmol) at 95 °C. The reaction was allowed to age for an additional 15 minutes and then cooled. The reaction mixture was diluted with water (50 ml) and extracted with ethyl acetate (3 x 50 ml). The organic extract was washed with 10% (m/v) aqueous LiCl solution (3 x 25 ml), dried over sodium sulfate, filtered and evaporated to give a residue that was flash chromatographed (15% EtOAc/hexane) to give 4-(difluoromethoxy)-3-methylbenzaldehyde (021GLM-053_1(2), 1.00 g, 5.37 mmol, 63%) as a yellow oil. This oil was combined with that of the previous experiment and passed through a Pasteur pipette column eluting with 10% EtOAC/hexane to give an oil that solidified on standing (1.315 g, 7.06 mmol, 67% over the two reactions). The proton NMR was consistent with the proposed structure.

**Synthesis of (4-(difluoromethoxy)-3-methylphenyl)(2-(3,5-difluorophenyl)-1,3-dithian-2-yl)methanol**

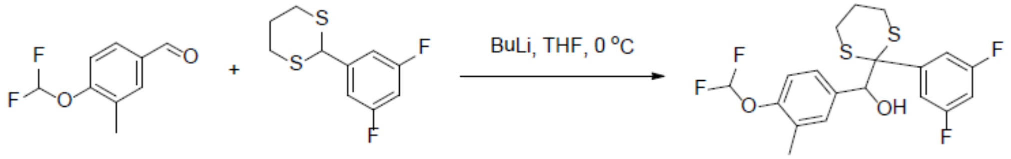


**Figure S16**. *Synthesis of (4-(difluoromethoxy)-3-methylphenyl)(2-(3,5-difluorophenyl)-1,3-dithian-2-yl)methanol*.

2-(3,5-Difluorophenyl)-1,3-dithiane (3.12 g, 13.43 mmol) was dissolved in 30 ml of dry THF and cooled to -10°C. BuLi (1.6M, 12.0 ml, 19.20 mmol) was added dropwise under nitrogen and the mixture was stirred for 15 min at -10°C to afford a blood-red solution. A solution of 4-(difluoromethoxy)-3-methylbenzaldehyde (2.50 g, 13.43 mmol) in THF (10 ml) was added dropwise and the mixture was stirred for 15 minutes, then warmed to ambient temperature over 10 minutes and quenched with saturated ammonium chloride solution. The organic phase was washed with brine and dried with sodium sulfate. The solvent was removed and the residue flash chromatographed (10%EtOAc/hexane) to give (4-(difluoromethoxy)-3-methylphenyl)(2-(3,5-difluorophenyl)-1,3-dithian-2-yl)methanol (3.07 g, 7.22 mmol, 55%) as a thick oil that solidified on standing. The NMR was consistent with the proposed structure.

**Synthesis of 2-(4-(difluoromethoxy)-3-methylphenyl)-1-(3,5-difluorophenyl)-2-hydroxyethanone**

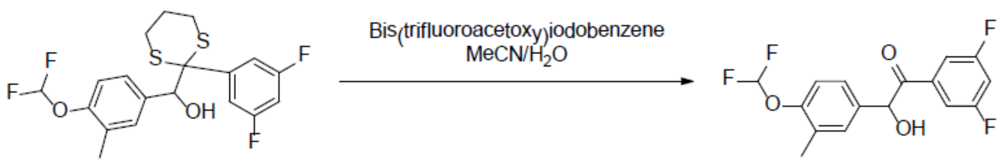


**Figure S17**. *Synthesis of 2-(4-(Difluoromethoxy)-3-methylphenyl)-1-(3,5-difluorophenyl)-2hydroxyethanone*.

(4-(Difluoromethoxy)-3-methylphenyl)(2-(3,5-difluorophenyl)-1,3-dithian-2-yl)methanol (3.07 g, 7.34 mmol) was dissolved in acetonitrile (15 ml) and water (2.5 ml). Bis(trifluoroacetoxy)iodobenzene (3.94 g, 9.17 mmol) in acetonitrile (10 ml) was slowly added at ambient temperature to the vigorously stirred solution. After 20 minutes TLC (20% EtOAc/hexane) analysis indicated a complete reaction. EtOAc (150 ml) was added and the mixture was rinsed with saturated sodium bicarbonate solution (50 ml) and brine (50 ml). The organic fractions were dried, and the solvent was removed in vacuo. The crude product was purified twice by flash column chromatography (10% EtOAc/hexane) to give 2-(4-(difluoromethoxy)-3-methylphenyl)-1-(3,5-difluorophenyl)-2-hydroxyethanone (0.853 g, 2.60 mmol, 35%) as a pale yellow oil. The proton NMR was consistent with the proposed structure.

**Synthesis of 1-(4-(difluoromethoxy)-3-methylphenyl)-2-(3,5-difluorophenyl)ethane-1,2-dione**

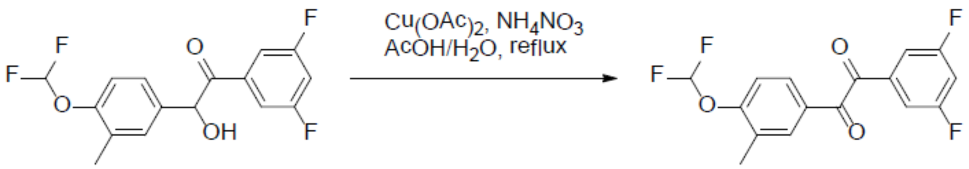


**Figure S18**. *Synthesis of 1-(4-(difluoromethoxy)-3-methylphenyl)-2-(3,5-difluorophenyl)ethane-1,2-dione*.

2-(4-(trifluoromethoxy)-3-methyl phenyl)-1-(3,5-fluorophenyl)-2-hydroxyethanone (0.853 g, 2.60 mmol) was dissolved in 80% acetic acid together with diacetoxycopper hydrate (52 mg, 0.26 mmol) and ammonium nitrate (0.156 g, 1.95 mmol). The mixture was refluxed for 90 minutes and then cooled. The reaction mixture was poured into ethyl acetate (50 ml) and washed with brine (2 x 25 ml), dried over sodium sulfate, filtered, and evaporated. The residue was azeotrope with toluene to remove acetic acid and the residue (0.737 g, 2.26 mmol, 87%) was used directly into the next stage.

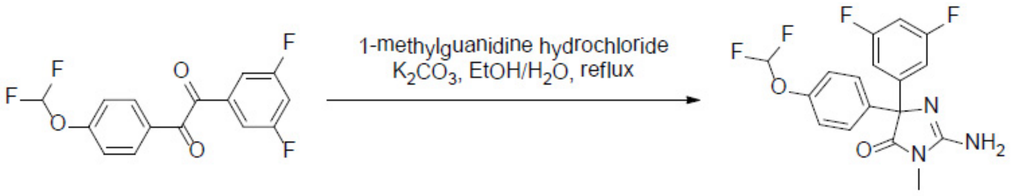


**Figure S19**. *Synthesis of 2-amino-4-(4-(difluoromethoxy)phenyl)-4-(3,5-difluorophenyl)-1-methyl-1H-imidazol-5(4H)-one*.

A mixture of 1-(4-(difluoromethoxy)-3-methylphenyl)-2-(3,5-difluorophenyl)ethane-1,2-dione (737 mg, 2.26 mmol) in ethanol (15 ml) and dioxane (15 ml) was added 1-methylguanidine hydrochloride (990 mg, 9.04 mmol) and stirred at ambient temperature for 15 minutes. Sodium carbonate (958 mg, 9.04 mmol) in water (5 ml) was added and the mixture was immersed into an oil bath at 85 °C and stirred for 3 hours. TLC (EtOAc) indicated a complete reaction. The reaction mixture was cooled to ambient temperature and concentrated. Purification by flash chromatography (EtOAc) afforded 2-amino-4-(4-(difluoromethoxy)-3-methyl phenyl)-4-(3,5-difluorophenyl)-1-methyl-1H-imidazol-5(4H)-one (0.52 g, 1.36 mmol, 60%) as a yellow solid. $^1$H NMR (400 MHz, d6-DMSO) δ 7.32-7.04 (m, 7H), 6.76 (s, 2H), 2.94 (s, 3H), 2.14 (s, 3H).; LC (260 nm): $R_t$ = 3.919 min, LC Purity: 96.1%, m/z (M+1): 382, LC (220 nm): $R_t$ = 3.922 min, %). LC Purity: 96.8 %.

*FAH3 Data analysis*

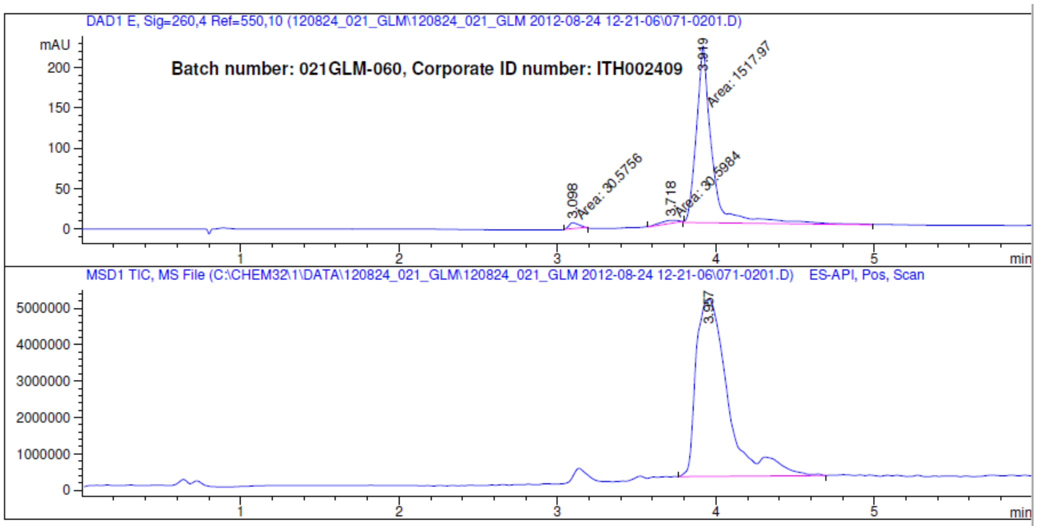


Area Percent Report

Sorted By : Signal
Multiplier: : 1.0000
Dilution: : 1.0000
Do not use Multiplier & Dilution Factor with ISTDs

Signal 1: DAD1 E, Sig=260,4 Ref=550,10

| Peak # | RetTime [min] | Type | Width [min] | Area [mAU*s] | Height [mAU] | Area % |
|---|---|---|---|---|---|---|
| 1 | 3.098 | MM | 0.0700 | 30.57557 | 7.27519 | 1.9362 |
| 2 | 3.718 | MM | 0.1364 | 30.59837 | 3.73874 | 1.9377 |
| 3 | 3.919 | MM | 0.1156 | 1517.96594 | 218.84267 | 96.1261 |

**Figure S20**. *LC: Chromatogram and Results for FAH3*

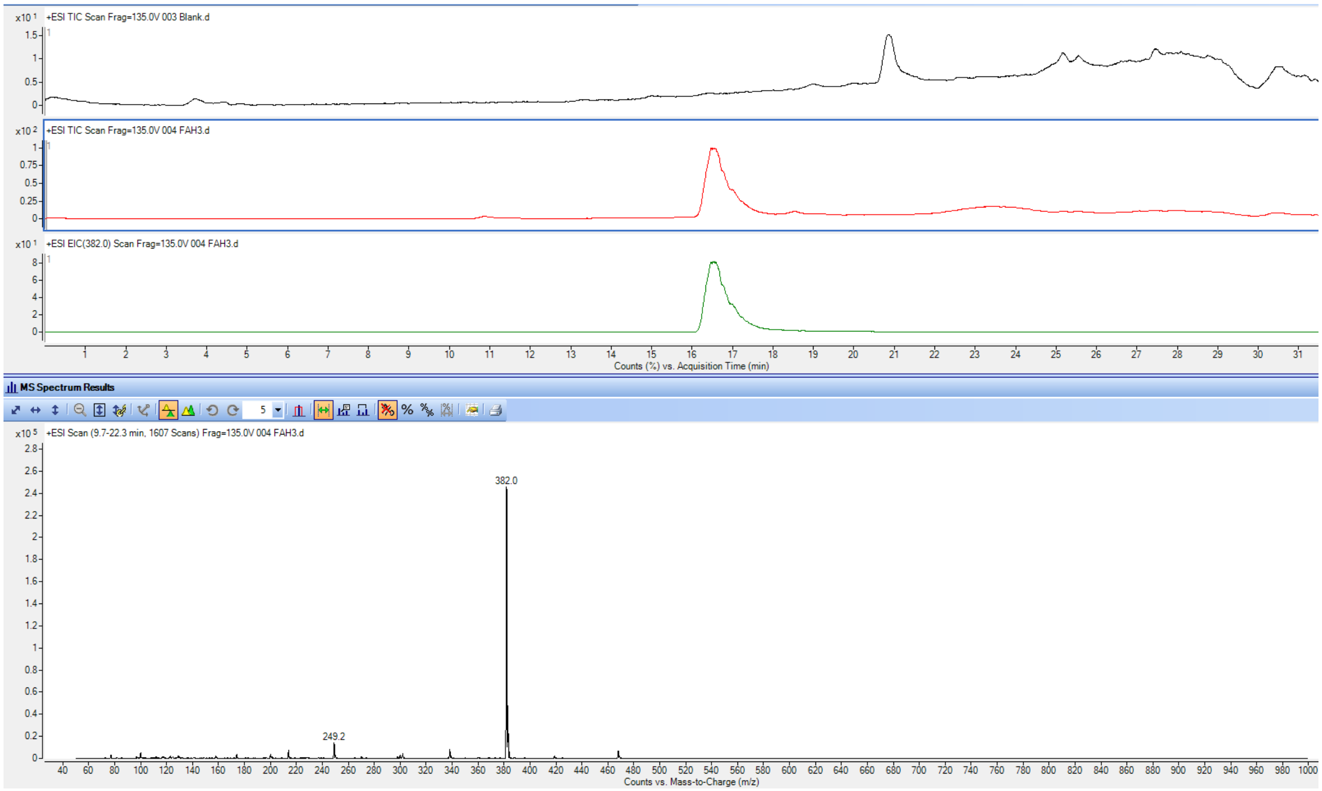


**Figure S21**. *LC-MS for FAH3. Blank TIC in black, FAH3 TIC in red, and FAH3 EIC in green*.

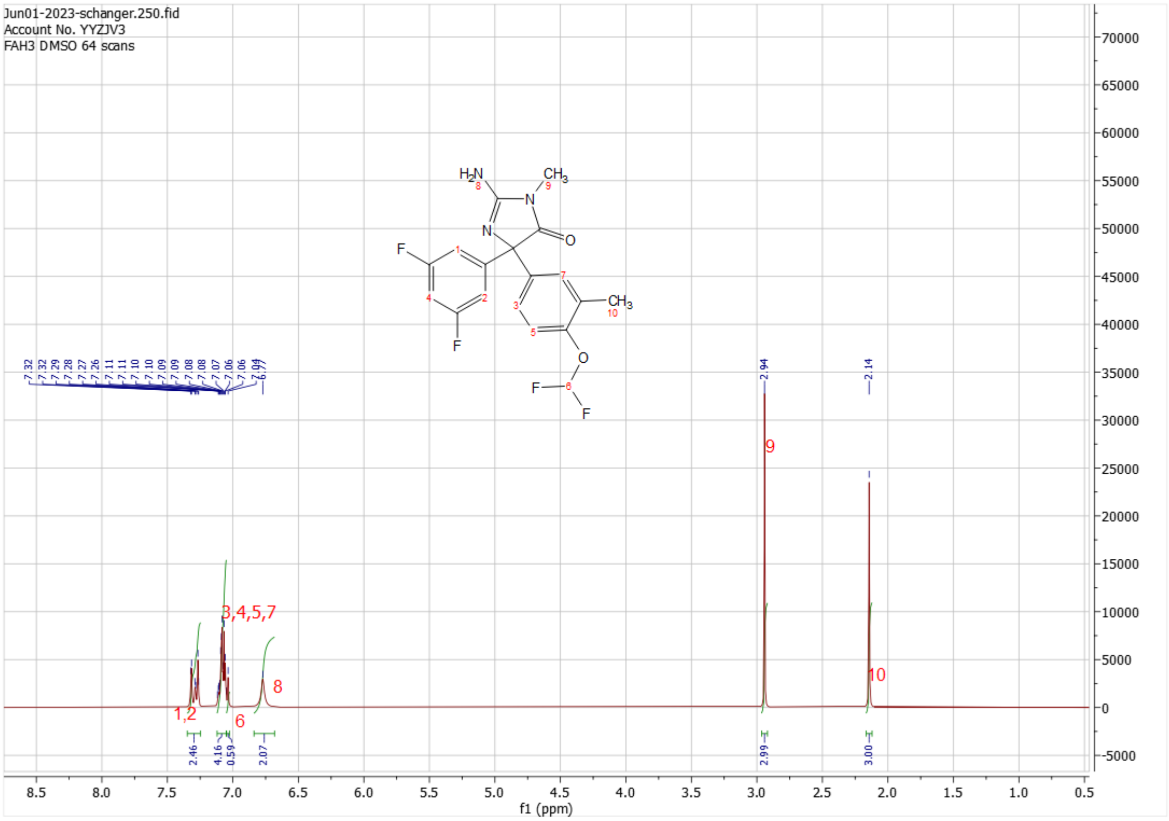


**Figure S22**. *NMR for FAH3*.

*FAH-17*

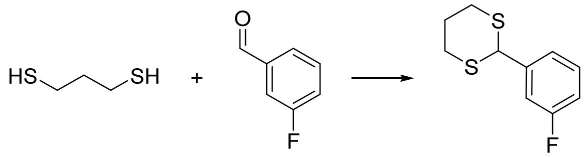

**Figure S23**. *Synthesis of 2-(3-fluorophenyl)-1,3-dithiane.*

$BF_3.OEt_2$ (4.30 ml, 34.8 mmol) was added dropwise to a solution of 1,3-propanedithiol (4.07 ml, 40.3 mmol) and 3-fluorobenzaldehyde (5.00 g, 40.3 mmol) in DCM (201 ml) at 0 °C. The reaction was stirred at ambient temperature for 1 hour where TLC (5% EtOAc/hexane) indicated a complete reaction. The reaction mixture was then diluted with DCM (50 ml), filtered through Celite (and the Celite pad was washed with additional DCM (3 x 50 ml)) and the filtrate washed with brine (100 ml), saturated $NaHCO_3$ (3 x 100 ml), 10% KOH solution (100 ml), water (100 ml) and brine (100 ml) and finally dried over sodium sulfate. The organic extract was filtered and evaporated. The product was purified using 5 % ethyl acetate:hexane to afford 2-(3-fluorophenyl)-1,3-dithiane ( 8.71 g, 39.0 mmol, 97 %) as an off-clear oil. The oil was used directly into the next step.

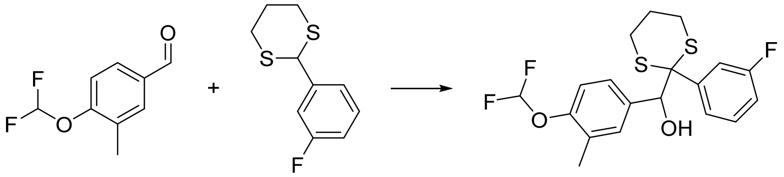

**Figure S24**. *Synthesis of (4-(difluoromethoxy)-3-methylphenyl)(2-(3-fluorophenyl)-1,3-dithian-2-yl)methanol.*

2-(3-fluorophenyl)-1,3-dithiane (4.00 g, 18.66 mmol) was dissolved in dry THF (93.5 mL) and cooled to -10 °C. nBuLi (1.6M, 14.00 ml, 22.40 mmol) was added dropwise under nitrogen and the mixture was stirred for 30 min at -10°C to afford a dark red solution. A solution of 4-(difluoromethoxy)-3-methylbenzaldehyde (3.47 g, 18.66 mmol) in THF (93.5 ml) was added dropwise and the mixture at -10 °C and was stirred for 15 minutes, then warmed to ambient temperature over 1 h and quenched with saturated ammonium chloride solution (7.5 ml) followed by dilution with EtOAc (50 ml). The organic phase was washed with water (2x20 ml), brine (1x20 ml) and dried with sodium sulfate. After filtration and concentration the crude product was purified by flash column chromatography (15% EtOAc/Hex) to give (4-(difluoromethoxy)-3-methylphenyl)(2-(3-fluorophenyl)-1,3-dithian-2-yl)methanol (6.00 g, 14.96 mmol, 80%) as an oil.

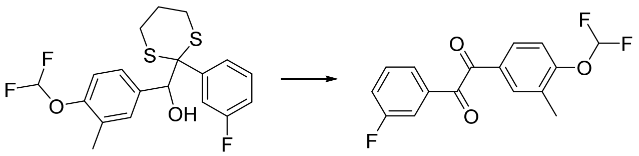

**Figure S25**. *Synthesis of 1-(4-(difluoromethoxy)-3-methylphenyl)-2-(3-fluorophenyl)ethane-1,2-dione.*

(4-(difluoromethoxy)-3-methylphenyl)(2-(3-fluorophenyl)-1,3-dithian-2-yl)methanol (5.55 g, 13.86 mmol) was dissolved in dichloromethane (175 ml) and tert-butanol (37.1 ml, 388 mmol) under nitrogen atmosphere. Dess-Martin Periodinane (14.69 g, 34.6 mmol) was added and the reaction was stirred overnight at room temperature. Sodium thiosulphate (5 ml, 1M) was added and the layers were separated. The organic phase was washed with sodium hydrogen carbonate and the solvent was evaporated. Purification on column chromatography in 5 % ethyl acetate hexane gave 1-(4-(difluoromethoxy)-3-methylphenyl)-2-(3-fluorophenyl)ethane-1,2-dione (3.12 g, 10.12 mmol, 73 %) as a yellow solid.

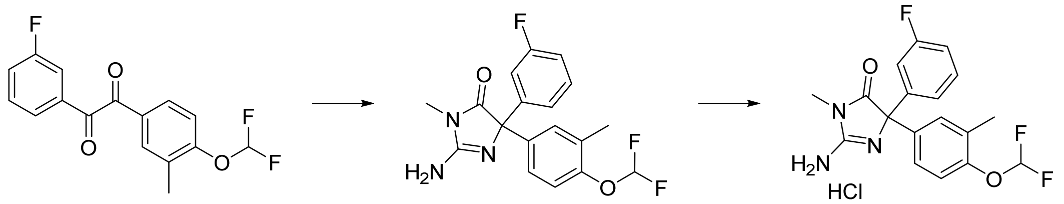


**Figure S26**. *Synthesis of 2-amino-4-(4-(difluoromethoxy)-3-methylphenyl)-4-(3-fluorophenyl)-1-methyl-1H-imidazol-5(4H)-one hydrochloride.*

Sodium carbonate (4.13 g, 38.9 mmol) in water (58.5 mL) was added into a mixture of 1-(4-(difluoromethoxy)-3-methylphenyl)-2-(3-fluorophenyl) ethane-1,2-dione (3.00 g, 9.73 mmol), 1-methylguanidine hydrochloride (4.27 g, 38.9 mmol), dioxane (151 mL), and ethyl alcohol (196mL). The reaction mixture was stirred at 85 °C for overnight. The volatiles were removed in vacuo, and the residue was taken in chloroform (50 ml) and washed with water (2 x 15 mL). The organic extracts were dried over $MgSO_4$. Evaporation and purification by column chromatography (50 - 90 % ethyl acetate: hexane) gave 2-amino-4-(4-(difluoromethoxy)-3-methylphenyl)-4-(3-fluorophenyl)-1-methyl-1H-imidazol-5(4H)-one (2.85 g, 7.53 mmol, 77 %) as an off-white solid. $^1$H NMR ($CDCl_3$): 7.32-7.18 (m, 5H), 6.99-6.96 (m, 2H), 6.48 (t, 1H, $J$ = 74.1 Hz), 5.77 (brs, 2H), 3.11 (s, 3H), 2.25 (s, 3H); $^{13}$C NMR ($CDCl_3$): 163.91, 161.47, 155.73, 149.22, 138.02, 130.01, 129.02, 125.74, 122.69, 118.76, 116.18, 114.74, 114.53, 114.37, 114.14, 113.60, 74.92, 25.86, 16.34 (please note: due to presence of fluorine atoms, $J^2_{C-F}$ – $J^4_{C-F}$ couplings giving rise to poorly resolved triplets and doublets are noted); LC (220 nm): $R_t$ = 3.85 min, purity 96%; MS: For $C_{18}H_{16}F_3N_3O_2$ expect M+H = 364.3 obtained 364.0

2-amino-4-(4-(difluoromethoxy)-3-methylphenyl)-4-(3-fluorophenyl)-1-methyl-1H-imidazol-5(4H)-one (0.50 g, 1.38 mmol) was dissolved in anhydrous DCM (66 ml) followed by addition of HCl (1M in diethyl ether, 2.2 ml). The mixture was stirred at room temperature for 5 min and the solvent evaporated in vacuo to yield 2-amino-4-(4-(difluoromethoxy)-3-methylphenyl)-4-(3-fluorophenyl)-1-methyl-1H-imidazol-5(4H)-one hydrochloride (0.50 g, 1.20 mmol, 87%) as a white solid. $^1$H NMR ($d_6$-DMSO): 11.78 (brs, 1H), 9.73 (brs, 1H), 7.53-7.06 (m, 8H), 3.19 (s, 3H), 2.23 (s, 3H); $^{13}$C NMR ($d_6$-DMSO): 176.62, 176.99, 172.57, 163.63, 161.20, 157.95, 150.07, 150.04, 140.36, 140.30, 134.40, 129.86, 126.60, 123.70, 119.52, 119.00, 116.96, 116.47, 116.26, 114.64, 114.41, 70.30, 27.43, 16.4 0(please note: due to presence of fluorine atoms, $J^2_{C-F}$ – $J^4_{C-F}$ couplings giving rise to poorly resolved triplets and doublets are noted); LC (220 nm): $R_t$ = 3.84 min, purity 96.5%; MS: For $C_{18}H_{16}F_3N_3O_2$ expect $[M+H]^+$ = 364.3 obtained 364.1

*Data analysis for FAH17*

```
Acq. Operator   : SIYANDA                        Seq. Line :    2
Acq. Instrument : LCQ                             Location : Vial 11
Injection Date  : 12/12/2013 11:10:48 AM               Inj :    1
                                                Inj Volume : 5.0 µl
Acq. Method     : Z:\DATA\131212_021_STM\131212A_021_STM 2013-12-12 11-01-05\220_POS_100-650MS.M
Last changed    : 1/30/2013 8:00:11 AM by MAMOALOSI
Analysis Method : C:\CHEM32\1\METHODS\260_POS_100-1000MS.M
Last changed    : 3/7/2013 3:44:16 PM by SUSAN
Method Info     :  260nm NEG 100-500MS
```

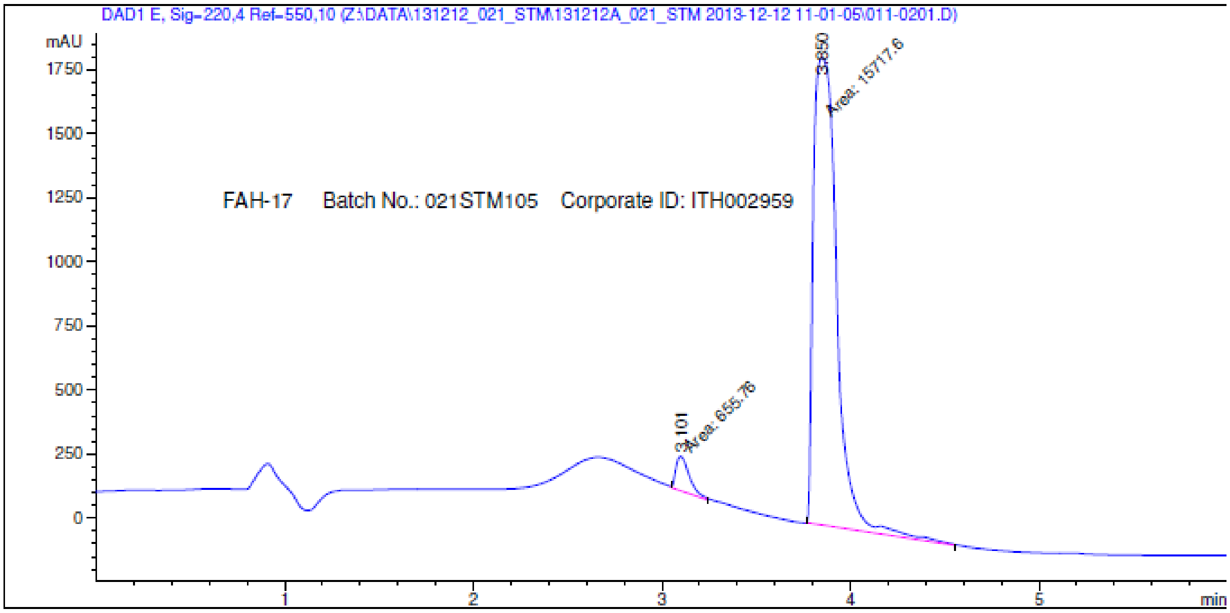


```
-----------------------------------------------------------------------
                         Area Percent Report
-----------------------------------------------------------------------

Sorted By              :        Signal
Multiplier:                     :        1.0000
Dilution:                       :        1.0000
Do not use Multiplier & Dilution Factor with ISTDs


Signal 1: DAD1 E, Sig=220,4 Ref=550,10

Peak RetTime Type  Width     Area      Height     Area
  #   [min]        [min]   [mAU*s]     [mAU]        %
----|-------|----|-------|----------|----------|--------|
   1   3.101 MM    0.0818  655.76044  133.53052   4.0050
   2   3.850 MM    0.1438 1.57176e4  1822.25830  95.9950

Totals :                  1.63734e4  1955.78882
```

**Figure S27**. *LC: Chromatogram and Results for FAH17*

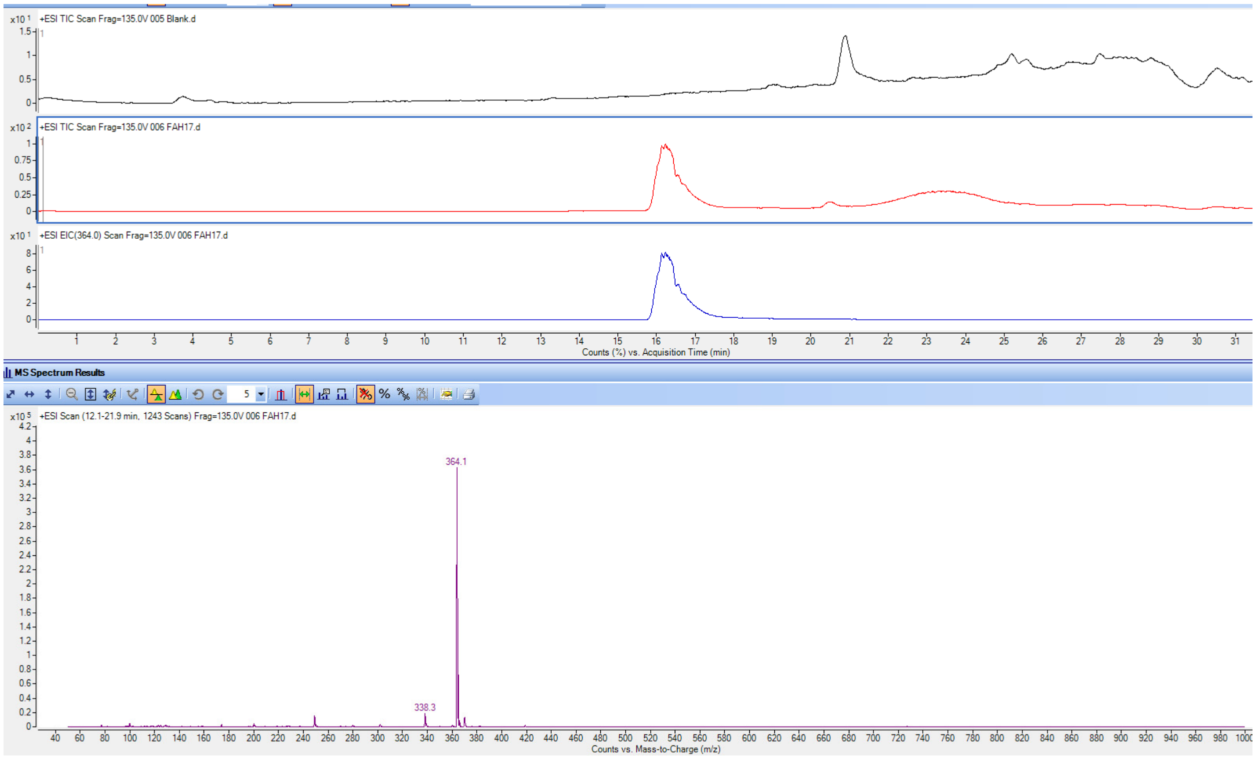


**Figure S28**. *LC-MS for FAH17. Blank TIC in black, FAH17 TIC in red, and FAH17 EIC in blue.*

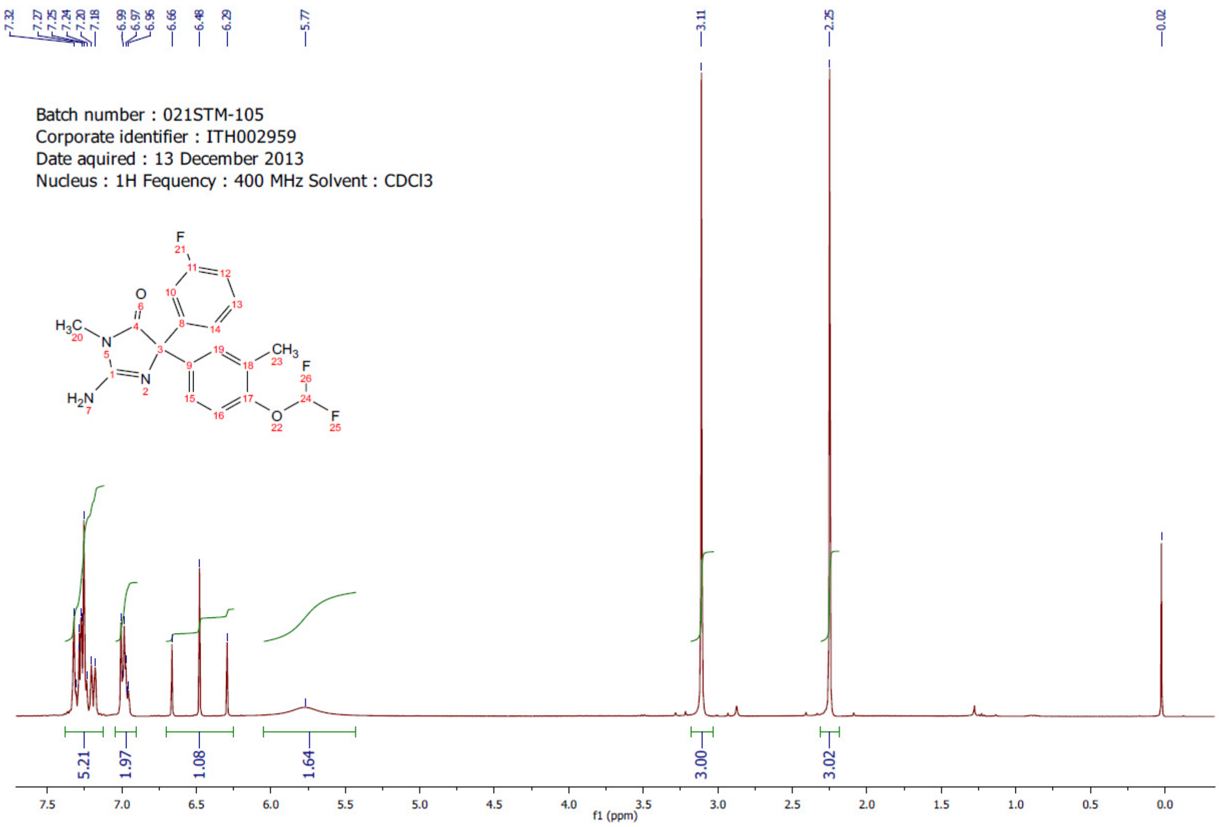


**Figure S29**. *NMR for FAH17*

*Synthetic Method of FAH17-HCl salt*

The preparation of FAH-17 HCl salt: 2-Amino-4-(4-(difluoromethoxy)-3-methylphenyl)-4-(3-fluorophenyl)-1-methyl-1H-imidazol-5(4H)-one (0.50 g, 1.38 mmol) was dissolved in anhydrous DCM (66 mL) followed by addition of HCl (1M in diethyl ether, 2.2 mL). The mixture was stirred at room temperature for 5 minutes and the solvent evaporated in vacuo to yield 2-amino-4-(4-(difluoromethoxy)-3-methylphenyl)-4-(3-fluorophenyl)-1-methyl-1H-imidazol-5(4H)-one hydrochloride (0.50 g, 1.21 mmol, 87%) as a white solid.

```
=====================================================================
Acq. Operator   : SIYANDA                          Seq. Line :    1
Acq. Instrument : LCQ                               Location : Vial 12
Injection Date  : 12/13/2013 9:35:26 AM                  Inj :    1
                                                 Inj Volume : 5.0 µl
Acq. Method     : Z:\DATA\131213_021_STM\131213_021_STM 2013-12-13 09-34-39\220_POS_100-650MS.M
Last changed    : 1/30/2013 8:00:11 AM by MAMOALOSI
Analysis Method : C:\CHEM32\1\METHODS\260_POS_100-1000MS.M
Last changed    : 3/7/2013 3:44:16 PM by SUSAN
Method Info     :  260nm NEG 100-500MS
```

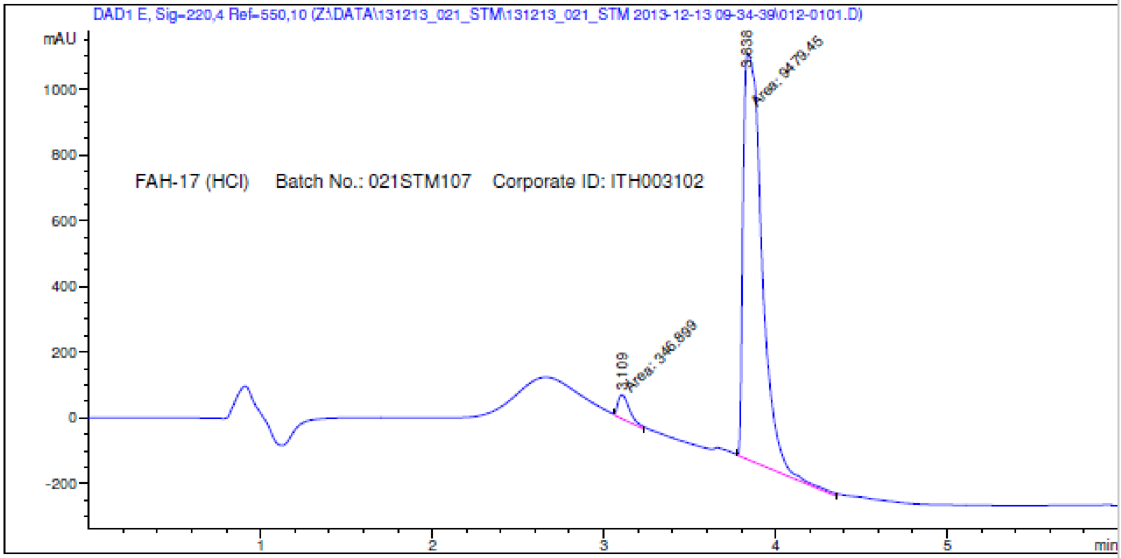


```
=====================================================================
                         Area Percent Report
=====================================================================

Sorted By             :      Signal
Multiplier:                  :       1.0000
Dilution:                    :       1.0000
Do not use Multiplier & Dilution Factor with ISTDs


Signal 1: DAD1 E, Sig=220,4 Ref=550,10

Peak RetTime Type  Width     Area      Height     Area
  #   [min]        [min]   [mAU*s]     [mAU]        %
----|-------|----|-------|----------|----------|--------|
   1   3.109 MM    0.0791  346.89920   73.07652    3.5303
   2   3.838 MM    0.1277 9479.45313 1236.86230   96.4697

Totals :                  9826.35233 1309.93882
```

**Figure S30**. *LC: Chromatogram and Results for FAH17 HCl salt*

```
Acq. Operator   : SIYANDA                          Seq. Line :    3
Acq. Instrument : LCQ                               Location : Vial 12
Injection Date  : 12/13/2013 9:53:43 AM                  Inj :    1
                                                  Inj Volume : 5.0 µl
Acq. Method     : Z:\DATA\131213_021_STM\131213_021_STM 2013-12-13 09-34-39\210_POS_100-650MS.M
Last changed    : 1/21/2012 10:02:08 AM by GLM
Analysis Method : C:\CHEM32\1\METHODS\260_POS_100-1000MS.M
Last changed    : 3/7/2013 3:44:16 PM by SUSAN
Method Info     :  260nm NEG 100-500MS
```

MS Spectrum

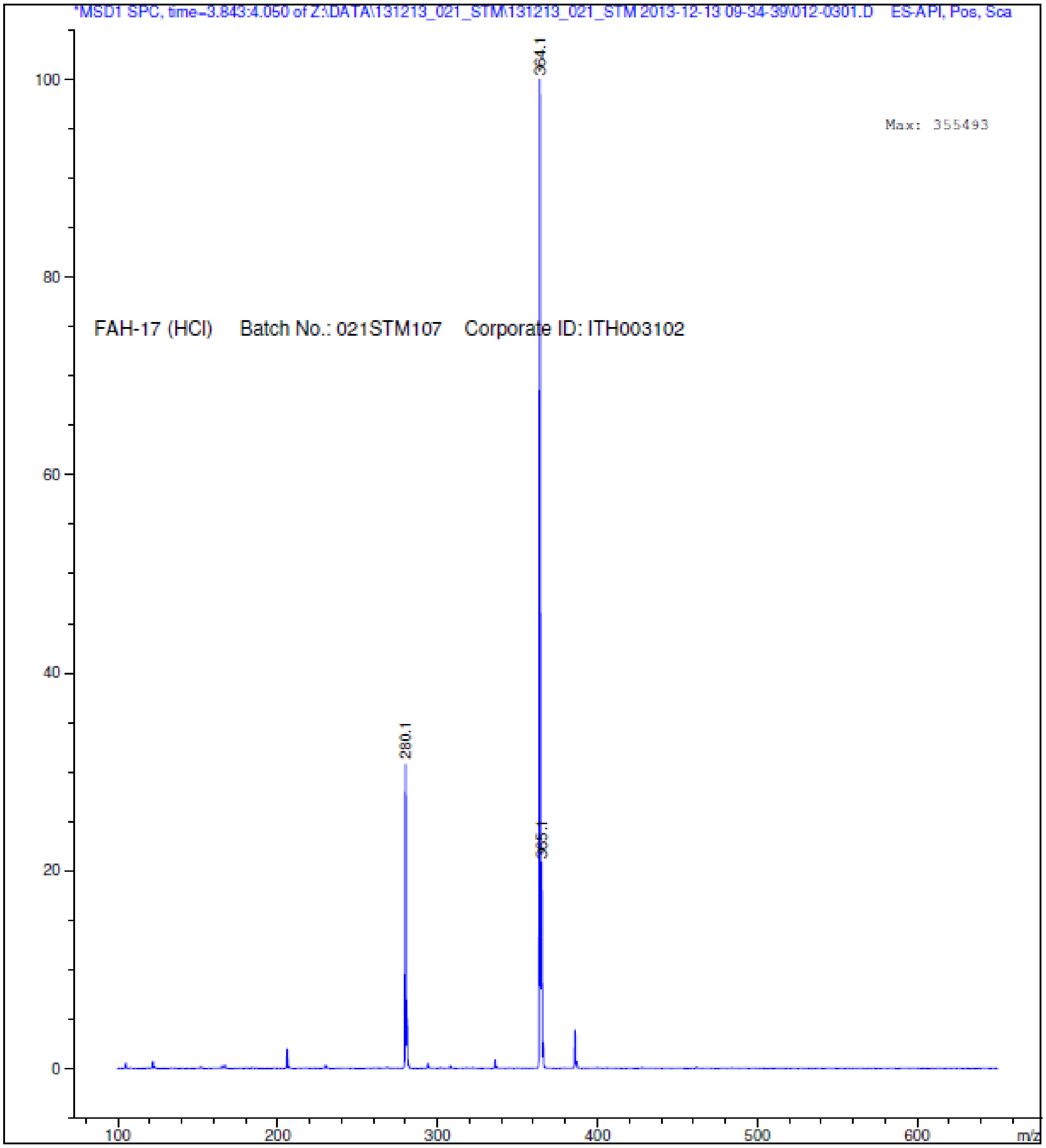


**Figure S31**. *Mass spectrum for FAH17 HCl salt*

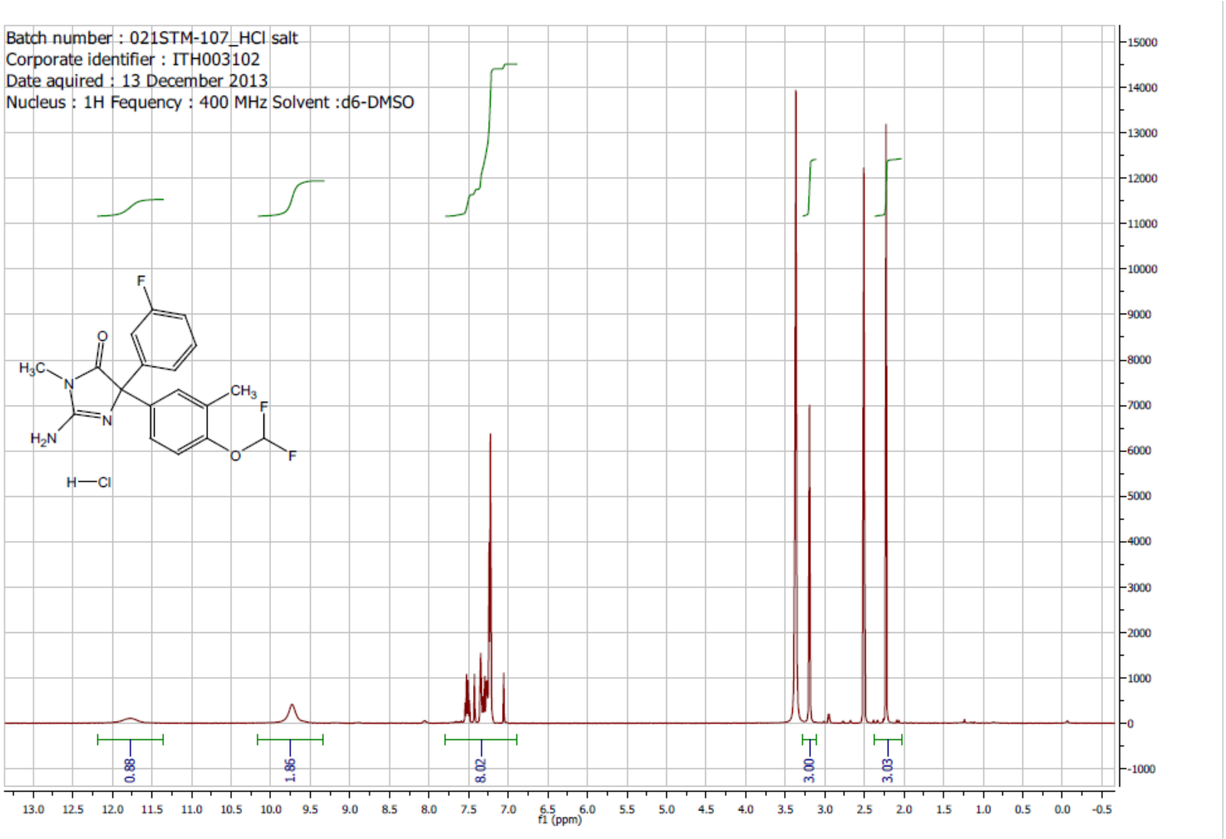


**Figure S32**. *H-NMR for FAH17 HCl salt*

*FAH65*

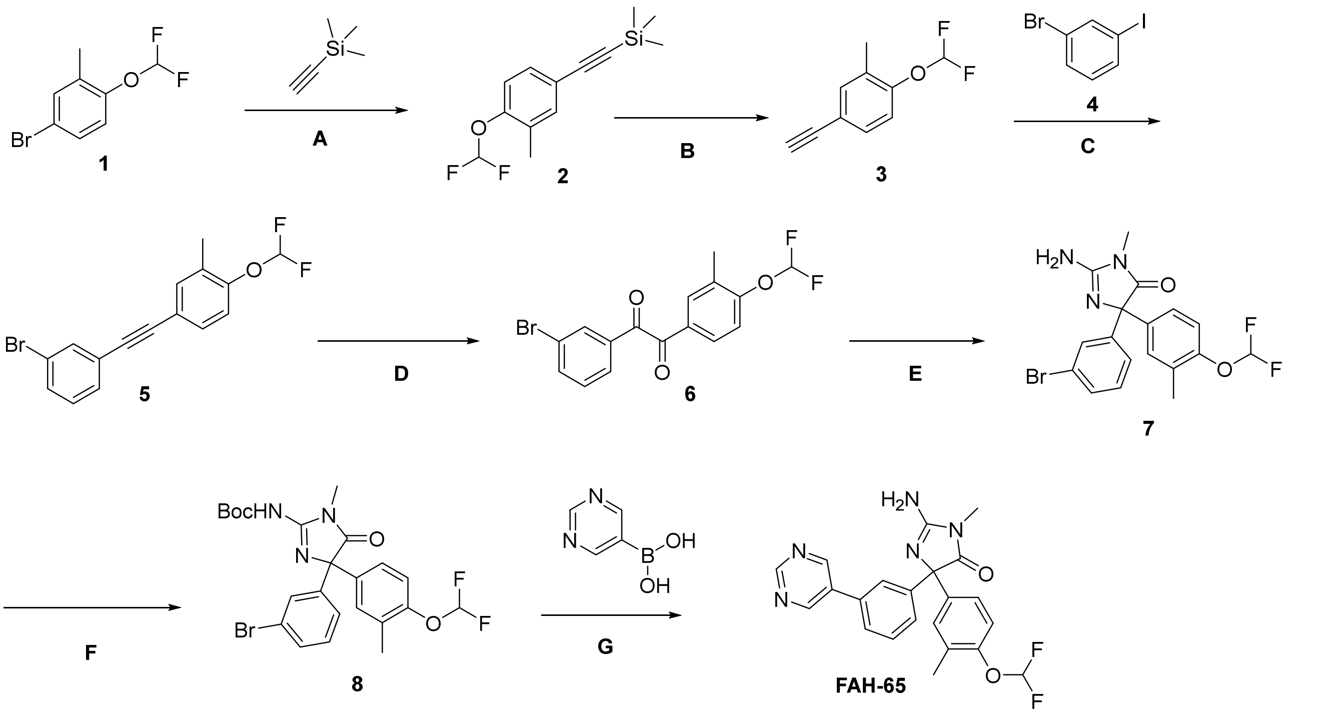

**Figure S33**. *Synthesis scheme for FAH65*

*General experimental methods and Protocols for FAH65*

$^{1}$H NMR spectra were recorded on an Oxford 400 MHz actively-shielded magnet with a Bruker AvanceIII HD Nanobay console, and a 5mm BBFO Z-gradient Smart Probe with Automatic Tune and Match or a Bruker 600 MHz Ultrashield Plus superconducting magnet with a Bruker Avance III console using a 5mm broad-band observe (BBO) probe with Automatic Tune and Match and a Z-gradient. Spectra are referenced on the residual solvent peak. LCMS data were acquired on an Advion CMS qaudropole mass spectrometer (pos/neg electrospray ionization) that was connected to a Gilson LC front-end. The data were analyzed with Advion's Data Express software.

**Synthesis of compound 2.** Dissolve 4-bromo-1-(difluoromethoxy)-2-methylbenzene **(1)** (10 g, 42.2 mmol) in dry DMF (100 mL). Trimethylsilylacetylene (7.01 mL, 50.6 mmol), and triethylamine (17.6 mL, 127 mmol) were added and the mixture was bubbled with argon gas for about 5 minutes. Copper(I)iodide (8.3 mg, 4.22 mmol) and bis(tpp)PdCl$_2$ (1.48 g, 2.11 mmol) were added and the mixture was stirred at 60 °C overnight. The reaction mixture was diluted with ether and washed with brine. The organic layer was separated, washed with water and dried using sodium sulfate, filtered and concentrated under reduced pressure. The crude mixture was purified using flash chromatography using hexane and ethyl acetate to yield light brown solid **(2)** (5.3 g, 49.4% yield).

**Synthesis of compound 3.** Dissolved ((4-(difluoromethoxy)-3-methylphenyl)ethynyl)trimethylsilane **(2)** (5.3 g, 20.8 mmol) in methanol (106 mL). Add potassium carbonate (7.2 g, 52.1 mmol) to the reaction mixture and stirred at room temperature overnight. The solids were filtered using vacuum filtration. The filtrate was diluted with 1:4 ethyl acetate:hexane and was washed with water. Water was back extracted with ethyl acetate/hexane. Organic layer was washed with brine, dried using sodium sulfate, filtered and the solvent was evaporated under reduced pressure to yield pale yellow solid **(3)** (3.5 g, 92.2% yield).

**Synthesis of compound 5.** A solution of diisopropylethyamine (20.1 mL, 115 mmol) and DMF (28 mL), argon gas was bubbled for about 15 minutes. 1-(difluoromethoxy)-4-ethynyl-2-methylbenzene **(3)** (3.5 g, 19.2 mmol) and 1-bromo-3-iodobenzene **(4)** (5.44 g, 19.2 mmol) were added to the reaction mixture and argon gas was bubbled for 5 minutes. Copper(I)iodide (366 mg, 1.92 mmol) and bis(tpp)$PdCl_2$ (674 mg, 961 μmol) were added to the reaction mixture and was stirred at 60 °C overnight. The mixture was diluted with ether and washed with water. The mixture was filtered over celite and the organic layer was separated. The aqueous layer was back extracted with ether. Combined organic layers were washed with saturated ammonium chloride solution, dried using sodium sulfate, filtered and the solvent was evaporated under reduced pressure. The crude mixture was partitioned with hexane and water. Solids were filtered, the organic layer was separated and dried using sodium sulfate. The mixture was filtered, and the solvent was evaporated under reduced pressure. The crude mixture was purified by passing through a column of silica gel using hexane as gradient to yield brown oil **(5)** (6.3 g, 97.3% yield).

**Synthesis of compound 6.** Dissolve 4-((3-bromophenyl)ethynyl)-1-(difluoromethoxy)-2-methylbenzene **(5)** (6.3 g, 18.7 mmol) in DCM (252 mL) and acetic acid (12.8 mL, 224 mmol) was added. Tetrabutylammonium bromide (9.04 g, 28 mmol) and potassium permanganate (8.86 g, 56.1 mmol) were added to the reaction mixture and was stirred at room temperature for 2 hours. The mixture was diluted with DCM and was quenched with 5% sodium metabisulphite. The organic layer was separated and dried using sodium sulfate. The crude mixture was purified using flash chromatography using hexane and ethyl acetate as gradients to yield pale yellow solid **(6)** (5.4 g, 78% yield).

**Synthesis of compound 7.** To a solution of 1-(3-bromophenyl)-2-(4-(difluoromethoxy)-3-methylphenyl)ethane-1,2-dione **(6)** (5.4 g, 14.6 mmol) in isopropanol (81 mL), add sodium carbonate (2.71 g, 25.6 mmol) and 1-methyl guanidine hydrochloride (2.4 g, 21.9 mmol) and was stirred at 80 °C for 2 hours. After completion of the reaction, the reaction mixture was diluted with isopropanol and chloroform and washed with water. The aqueous phase was back extracted with chloroform. The combined organic layers were dried using sodium sulfate, filtered and concentrated under reduced pressure to yield colorless solid **(7)** (6.1 g, 98.3%).

**Synthesis of compound 8.** Dissolve 2-amino-5-(3-bromophenyl)-5-(4-(difluoromethoxy)-3-methylphenyl)-3-methyl-3,5-dihydro-4H-imidazol-4-one **(7)** (6.1 g, 14.4 mmol) in DCM (61 mL) and cool it to 0 C. Di-tert-butyl-dicarbonate (3.29 g, 15.1 mmol) was added to the reaction mixture and was stirred at room temperature overnight. After completion of the reaction, the reaction mixture was concentrated under reduced pressure. The crude mixture was purified using flash chromatography using hexane and ethyl acetate as gradients to yield off-white solid **(8)** (3.5 g, 46.4 %).

**Synthesis of 2-amino-5-(4-(difluoromethoxy)-3-methylphenyl)-3-methyl-5-(3-(pyrimidin-5-yl)phenyl)-3,5-dihydro-4H-imidazol-4-one (FAH-65).** Dissolved tert-butyl (4-(3-bromophenyl)-4-(4-(difluoromethoxy)-3-methylphenyl)-1-methyl-5-oxo-4,5-dihydro-1H-imidazol-2-yl)carbamate **(8)** (2.0 g, 3.81 mmol) in dioxane (10 mL). Pyrimidine-5-boronic acid (473 mg, 3.81 mmol), 2 M potassium carbonate solution (5.82 mL) and Xphos Palladacycle Gen.2 (300 mg, 381 μmol) were added and the mixture was heated at 110 °C for 2 h in a $N_2$ flushed sealed vial on a hot plate. The reaction mixture was partitioned with water and chloroform and further extracted to obtain the product. The combined organic layers were washed with brine, dried using sodium sulfate, and concentrated under reduced pressure. On an 80 g silica column, the crude was loaded and eluted with 0-30% ACN/DCM (10 CV) then 2 CV at 30% to get the Boc-protected compound. After concentrating down, the fraction was dissolved in DCM and added 1 g of charcoal, stirred for 10 min, filtered through a Celite plug and concentrated to obtain a pale-yellow solid (1.4 g). Then continued to elute the column with 10% MeOH/DCM to get de-Boc product. After concentrating down, the fraction was dissolved in MeOH/DCM and added charcoal, stirred for 10 min, filtered through a Celite plug and concentrated to obtain 350 mg of the title compound.

The Boc-product (1.4 g) was dissolved in DCM and TFA was added. The mixture was stirred at 40 °C for 2 h. The reaction was concentrated, and partitioned in between EA and $Na_2CO_3$ solution. The organic layer was washed with brine, dried over MgSO4, concentrated, and triturated with DCM to give an off-white solid (820 mg). The multiple crops of product were combined in acetonitrile and DCM in 20 ml vial and concentrated to obtain a uniform solid for characterization. LCMS (+esi) M+H+=424.4.

$^1$H NMR (600 MHz, METHANOL-$d_4$) δ = 9.12 (s, 1H), 9.00 (s, 2H), 7.69 (s, 1H), 7.68 - 7.65 (m, 1H), 7.55 - 7.50 (m, 2H), 7.28 (d, $J$ = 1.7 Hz, 1H), 7.24 (dd, $J$ = 2.1, 8.5 Hz, 1H), 7.07 (d, $J$ = 8.5 Hz, 1H), 6.91 - 6.64 (m, 1H), 3.15 (s, 3H), 2.23 (s, 3H).

*Data analysis for FAH65*

| | | | |
|---|---|---|---|
| *Acquisition Time (sec)* | 2.7263 | | |
| *Comment* | AV-600 ZBO proton starting parameters 11/16/08 RN | | |
| *D* | 0.1 | *D1* | 0.1 |
| *DE* | 6 | *DS* | 0 |
| *Date* | 06 Dec 2018 12:38:49 | | |
| *Date Stamp* | 06 Dec 2018 12:38:49 | | |
| *File Name* | \\10.1.100.200\chemgroup\HNMR\jasdug_NMR\JD-FAH65\4\PDATA\1\1r | | |
| *Frequency (MHz)* | 600.1300 | *GB* | 0 |
| *INSTRUM* | <AV-600> | *LB* | 0.3 |
| *NS* | 16 | *Nucleus* | 1H |
| *Number of Transients* | 16 | *Origin* | AV-600 |
| *Original Points Count* | 32768 | *Owner* | user |
| *PC* | 4 | | |
| *PROBHD* | <5 mm PABBO BB-1H/D Z-GRD Z847801/0074 > | | |
| *PULPROG* | <zg30> | *Points Count* | 131072 |
| *Pulse Sequence* | zg30 | *Receiver Gain* | 287.00 |
| *SF* | 600.130017072875 | | |
| *SFO1* | 600.133900845 | | |
| *SI* | 131072 | *SSB* | 0 |
| *SW(cyclical) (Hz)* | 12019.23 | *SWH* | 12019.2307692308 |
| *Solvent* | METHANOL-d4 | | |
| *Spectrum Offset (Hz)* | 3886.9551 | *Spectrum Type* | standard |
| *Sweep Width (Hz)* | 12019.14 | *TD* | 65536 |
| *TD0* | 2 | *TE* | 298 |
| *Temperature (degree C)* | 25.000 | *UNC1* | <1H> |
| *WDW* | 1 | | |

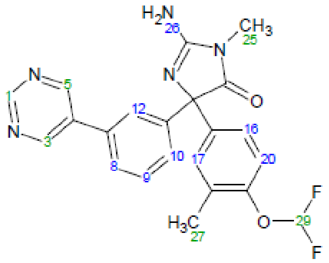


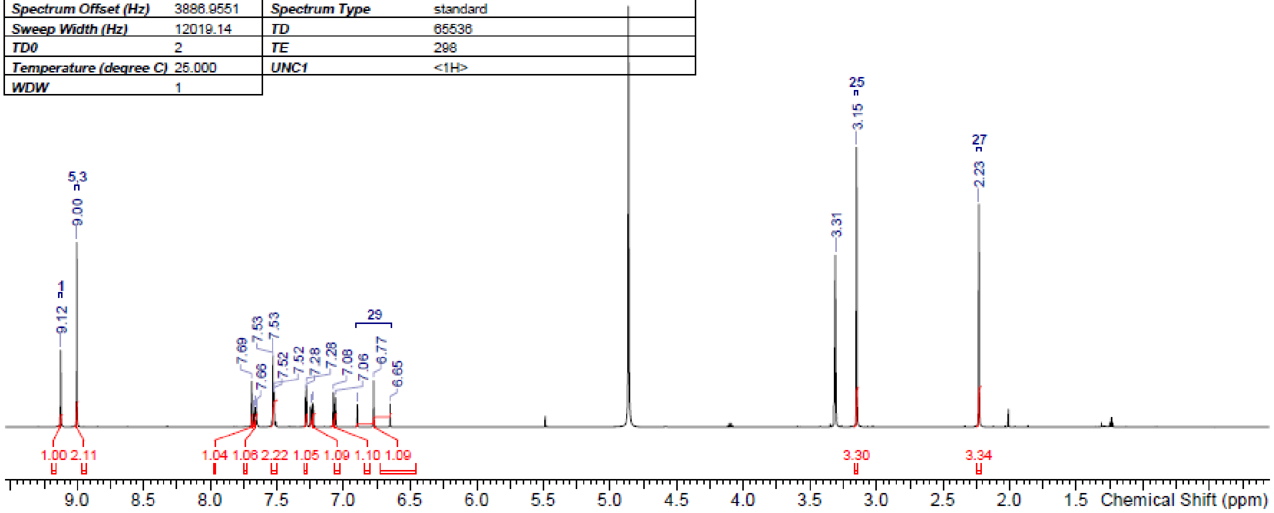


**Figure S34.** *NMR spectra FAH65*

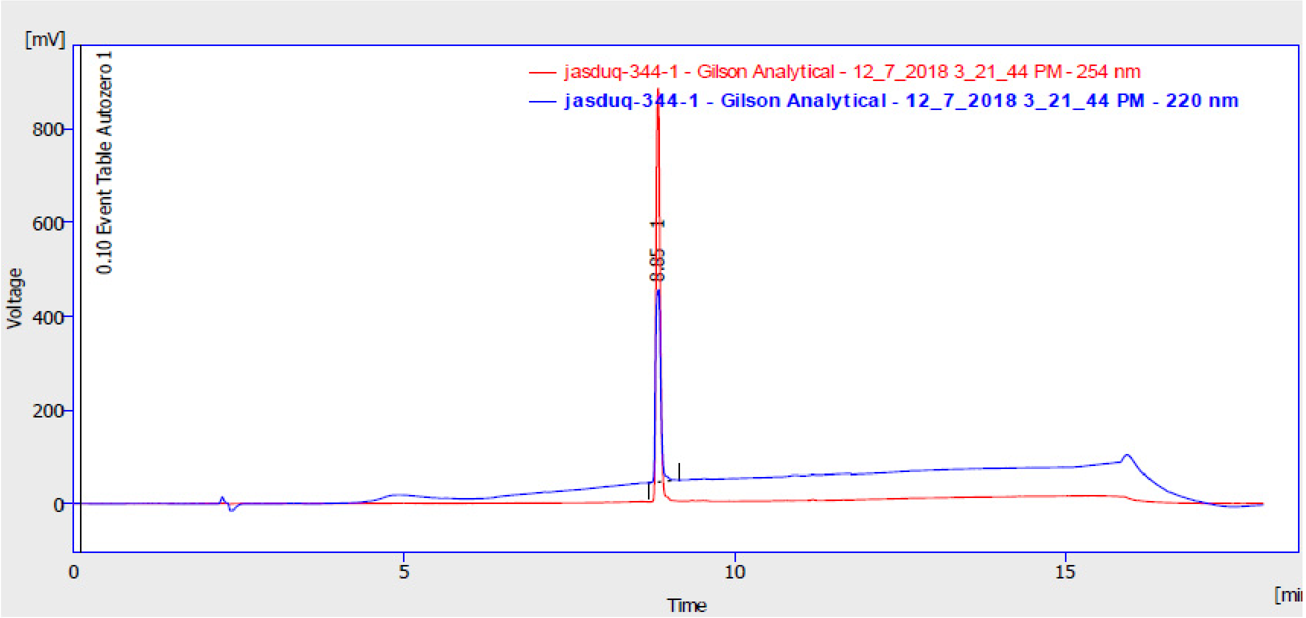


*Result Table (Uncal - jasduq-344-1 - Gilson Analytical - 12_7_2018 3_21_44 PM - 254 nm)*

| | Reten. Time [min] | Area [mV.s] | Height [mV] | Area [%] | W05 [min] | Compound Name |
|---|---|---|---|---|---|---|
| 1 | 8.843 | 3841.528 | 883.091 | 100.0 | 0.07 | |
| | Total | 3841.528 | 883.091 | 100.0 | | |

*Result Table (Uncal - jasduq-344-1 - Gilson Analytical - 12_7_2018 3_21_44 PM - 220 nm)*

| | Reten. Time [min] | Area [mV.s] | Height [mV] | Area [%] | W05 [min] | Compound Name |
|---|---|---|---|---|---|---|
| 1 | 8.850 | 2253.120 | 409.369 | 100.0 | 0.09 | |
| | Total | 2253.120 | 409.369 | 100.0 | | |

**Figure S35.** *FAH65 HPLC data*

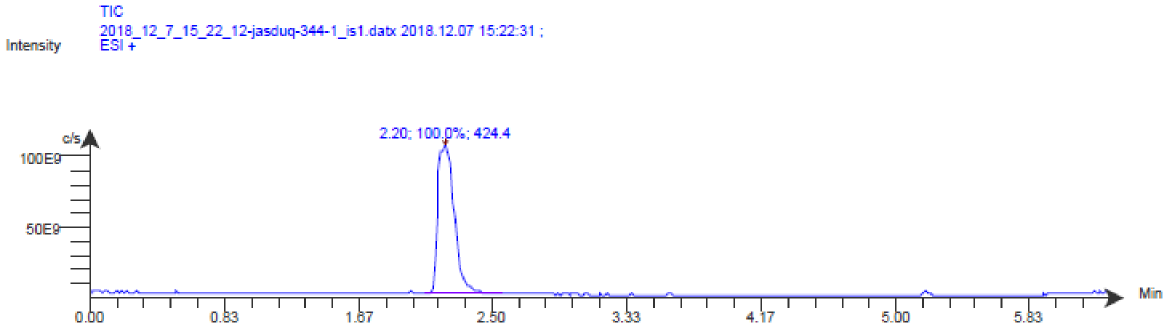


| Time (Peak Maximum M:S/Minutes) | Maximum Intensity (c/s) | Time (Peak Centroid M:S/Minutes) | Peak Area | % Peak Area | Peak Resolution | Base Peak Mass (m/z) | Label |
|---|---|---|---|---|---|---|---|
| 2.20 | 1.1E11 | 2.21 | 7.3E11 | 100.0 | 6.8 | 424.4 | |

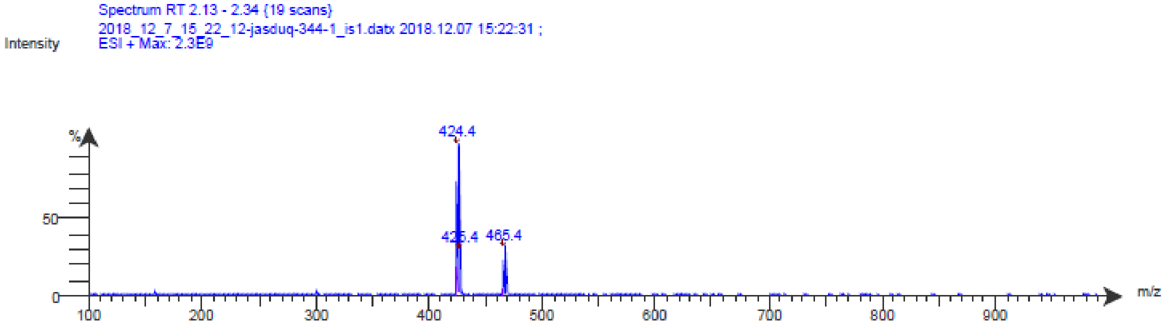


**Figure S36.** *FAH65 MS Data*

*FAH65 Enantiomer HCl salts*

*FAH65E(+)*

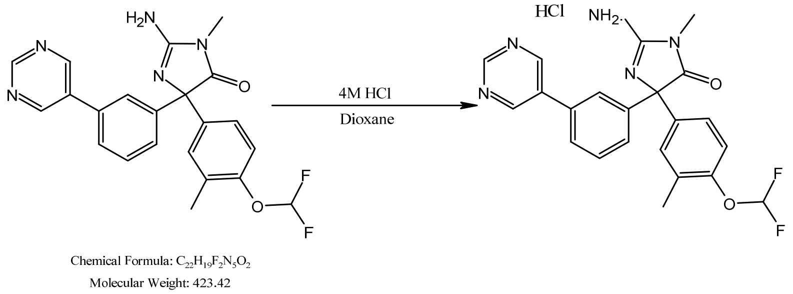


**Figure S37.** *Scheme for synthesis of FAH 65 P2 (E+) (100 mg batch) HCl salt (inactive compound)*

| Name | Molar Mass (g/mol) | Amount | Amount (mmol) | Eq |
|---|---|---|---|---|
| FAH 65 P2 (E+) | 423.42 | 90 mg | 0.213 | 1 |
| 4M HCl in 1,4-Dioxane | 5mL | | | |

A solution of FAH65 P2 (E+) (90mg, 0.213 mmol) in 4M HCl in dioxane (5 mL) was stirred for 6h at RT. After completion of the reaction, as indicated by TLC, dioxane was removed under vacuum and the resulting slurry was added to diethyl ether to precipitate a solid. Precipitates were filtered and washed three times with diethyl ether and dried overnight under a vacuum to afford FAH65 P2 (E+) as HCl salt (80.5mg, 82.3%, white solid)

*Data analysis for FAH65E(+)*

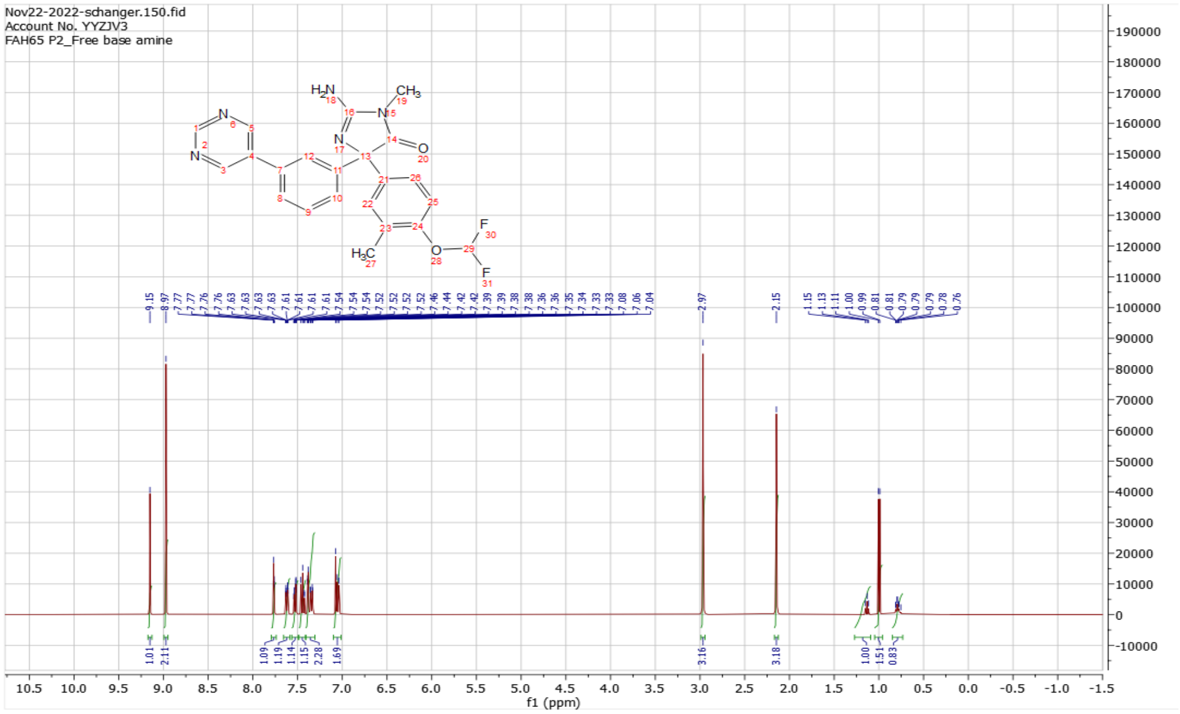


**Figure S38.** *A 64-scan proton NMR ($^1$H-NMR) for FAH65 P2 (E+)_free amine base utilizing default parameters*

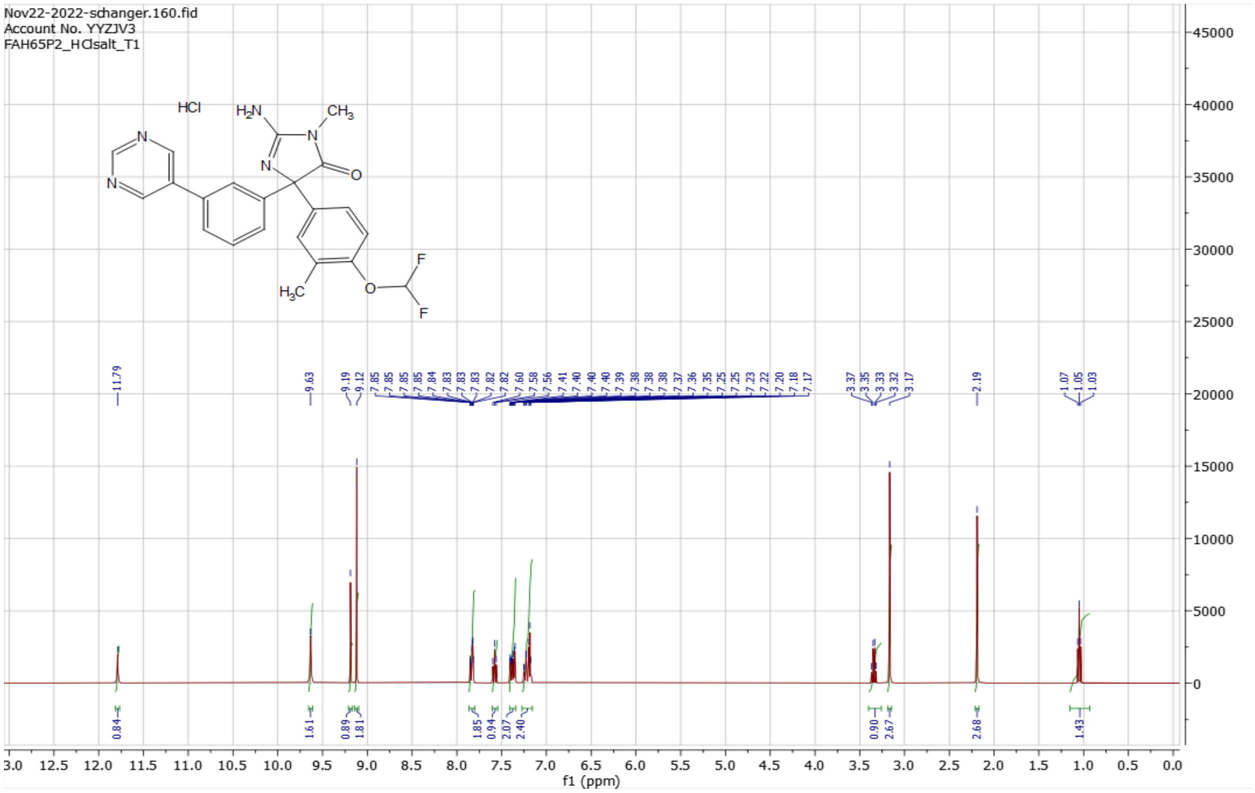


**Figure S39**. *A 64-scan proton NMR (¹H-NMR) for FAH65E(+) HCl salt (T1) utilizing default parameter*

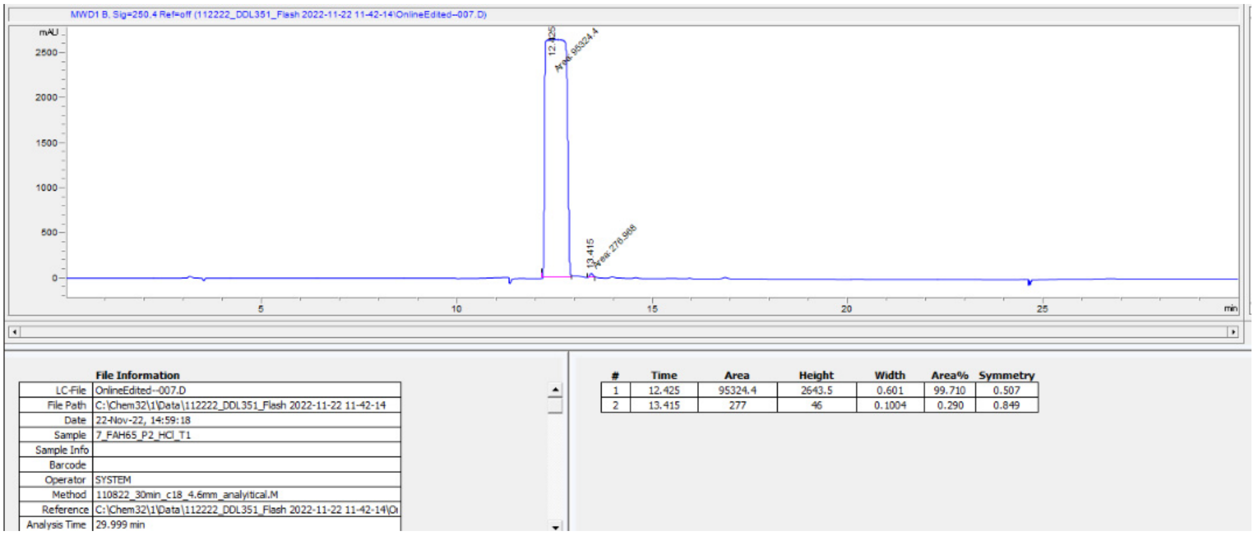


**Figure S40**. *HPLC-UVvis: Chromatogram and Results for FAH65E(+) HCl salt*

*Solubility of FAH65E(+)*

- FAH65E(+) HCl salt (equals to 20 mg/mL or greater than 20 mg/mL in water)
- FAH65E(+) Free amine base (less than 1 mg/mL in water)

*FAH65E(-)*

| Name | Molar Mass (g/mol) | Amount | Amount (mmol) | Eq |
|---|---|---|---|---|
| FAH65 P1 (E-) | 423.42 | 133.15 mg | 0.314 | 1 |
| 4M HCl in 1,4-Dioxane | 5mL | | | |

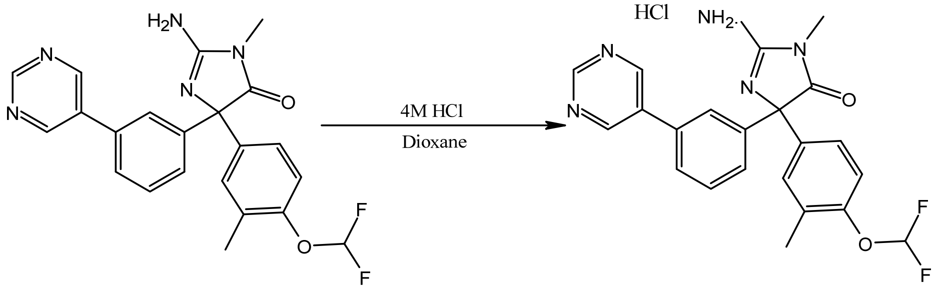


Chemical Formula: $C_{22}H_{19}F_2N_5O_2$
Molecular Weight: 423.42

**Figure S41.** *Synthesis scheme for FAH65E(-) (100 mg batch) HCl salt (active compound)*

A solution of FAH65E(-) (133.15 mg, 0.314 mmol) in 4M HCl in dioxane (5 mL) was stirred for 21h at RT. After completion of the reaction, as indicated by TLC, dioxane was removed under vacuum and the resulting slurry was added to diethyl ether to precipitate a solid. Precipitates were filtered and washed three times with diethyl ether and dried overnight under a vacuum to afford FAH65 P2 (E+) as HCl salt (118.32mg, 82%, white solid)

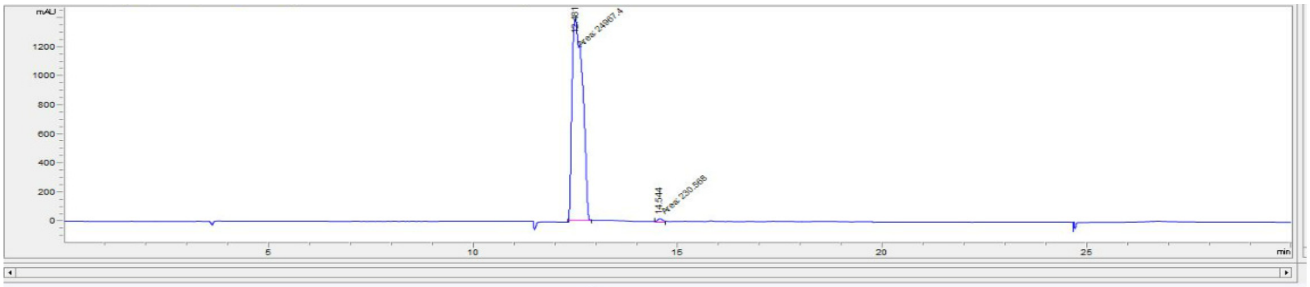


File Information

| | |
|---|---|
| LC-File | OnlineEdited--008.D |
| File Path | C:\Chem32\1\Data\120722_DDL10-SU_fractions 2022-12-07 12-44 |
| Date | 07-Dec-22, 16:34:35 |
| Sample | 7_FAH65P1_HCl_T2 |
| Sample Info | |
| Barcode | |
| Operator | SYSTEM |
| Method | 110822_30min_c18_4.6mm_analyitical.M |

| # | Time | Area | Height | Width | Area% | Symmetry |
|---|---|---|---|---|---|---|
| 1 | 12.481 | 24967.4 | 1397.6 | 0.2977 | 99.085 | 0.494 |
| 2 | 14.544 | 230.6 | 24.3 | 0.1582 | 0.915 | 0.756 |

**Figure S42**. *HPLC-UVvis: Chromatogram and Results for FAH65E(-) HCl salt*

Solubility of FAH65E(-) HCl

- FAH65E(-) HCl salt (100 mg batch) (equals to 20 mg/mL or greater than 20 mg/mL in water)
- FAH65E(-) Free amine base (less than 1 mg/mL in water)
